# Intercellular telomere transfer extends T cell lifespan

**DOI:** 10.1101/2020.10.09.331918

**Authors:** Bruno Vaz, Claudia Vuotto, Salvatore Valvo, Clara D’Ambra, Francesco Maria Esposito, Valerio Chiurchiù, Oliver Devine, Massimo Sanchez, Giovanna Borsellino, Derek Gilroy, Arne N. Akbar, Michael L. Dustin, Michael Karin, Alessio Lanna

**Affiliations:** Laboratory of NeuroImmuneSenescence, IRCCS Fondazione Santa Lucia, Rome. ITALY; Sentcell ltd, London, UK.; Experimental Neuroscience, IRCCS Fondazione Santa Lucia, Rome. ITALY; Kennedy Institute of Rheumatology, Nuffield Department of Orthopaedics, Rheumatology and Musculoskeletal Sciences, University of Oxford, Oxford. United Kingdom.; Institute of Translational Pharmacology, National Research Council, Rome. Italy.; Laboratory of resolution of NeuroInflammation, IRCCS Santa Lucia Foundation, Rome, Italy.; Division of Infection and Immunity, University College London, London, UK.; ISS, Core facilities, Rome. Italy.; NeuroImmunology Unit, IRCCS Fondazione Santa Lucia, Rome. ITALY; Division of Medicine, University College London, London, London, UK.; Laboratory of Gene regulation and Signal Transduction, University California San Diego. USA.

## Abstract

The common view is that T-lymphocytes activate telomerase, a DNA polymerase that extends telomeres at chromosome ends, to delay senescence. We show that independently of telomerase, T cells elongate telomeres by acquiring telomere vesicles from antigen-presenting cells (APCs). Upon contact with T cells, APCs degraded shelterin to donate telomeres, which were cleaved by TZAP, and then transferred in extracellular vesicles (EVs) at the immunological synapse. Telomere vesicles retained the Rad51 recombination factor that enabled them to fuse with T cell chromosomal ends causing an average lengthening of ∼3000 base pairs. Thus, we identify a previously unknown telomere transfer program that supports T cell lifespan.

---

Telomeres are TTAGGG repeats that protect chromosome ends and promote cellular lifespan^1^. In cells with short telomeres (<4kb), proliferative activity ceases and replicative senescence occurs^2^. Telomere shortening is observed in age-related pathologies and has provided a common mechanism linking senescence, cancer and ageing^3^. Although cells prevent telomere shortening by telomerase-dependent and independent pathways^4, 5^, it is not known whether telomeres can be transferred between cells as part of a telomere maintenance program. We found that T-lymphocytes, the largest subset of white blood cells that preserve telomeres to maintain life-long immunity^6^, elongate telomeres by acquiring telomeres in extracellular vesicles from APCs.

We conjugated primary human CD27^+^ CD28^+^ CD3^+^ CD4^+^ T cells with autologous APCs (peripheral blood mononuclear cells depleted of CD3^+^ cells) pulsed with cognate antigens (HBV, influenza and CMV lysates) to form antigen-specific synapses among the bulk population of T cells. Twenty-four hours later, we mechanically disrupted synapses and isolated T cell and APC fractions using antibodies to the major T-cell receptor (TCR) subunit CD3. Telomere-restriction fragment (TRF) analysis demonstrated telomere elongation up to 3 kb in the T cells and a concurrent telomere shortening in APCs after synapse formation (**Fig. 1a**). Increased telomere signal in T cells and a concomitant decrease in APCs were observed even in intact immunological synapses studied by immunofluorescence *in-situ* hybridization (IF-FISH, **Extended Data Fig. 1a, b**). Similar results were obtained by quantitative polymerase chain reaction (qPCR) with telomere specific primers (**Extended Data Fig. 1b**). No change in telomere length could be detected in immunological conjugates formed in the absence of antigens.

**Figure 1.**
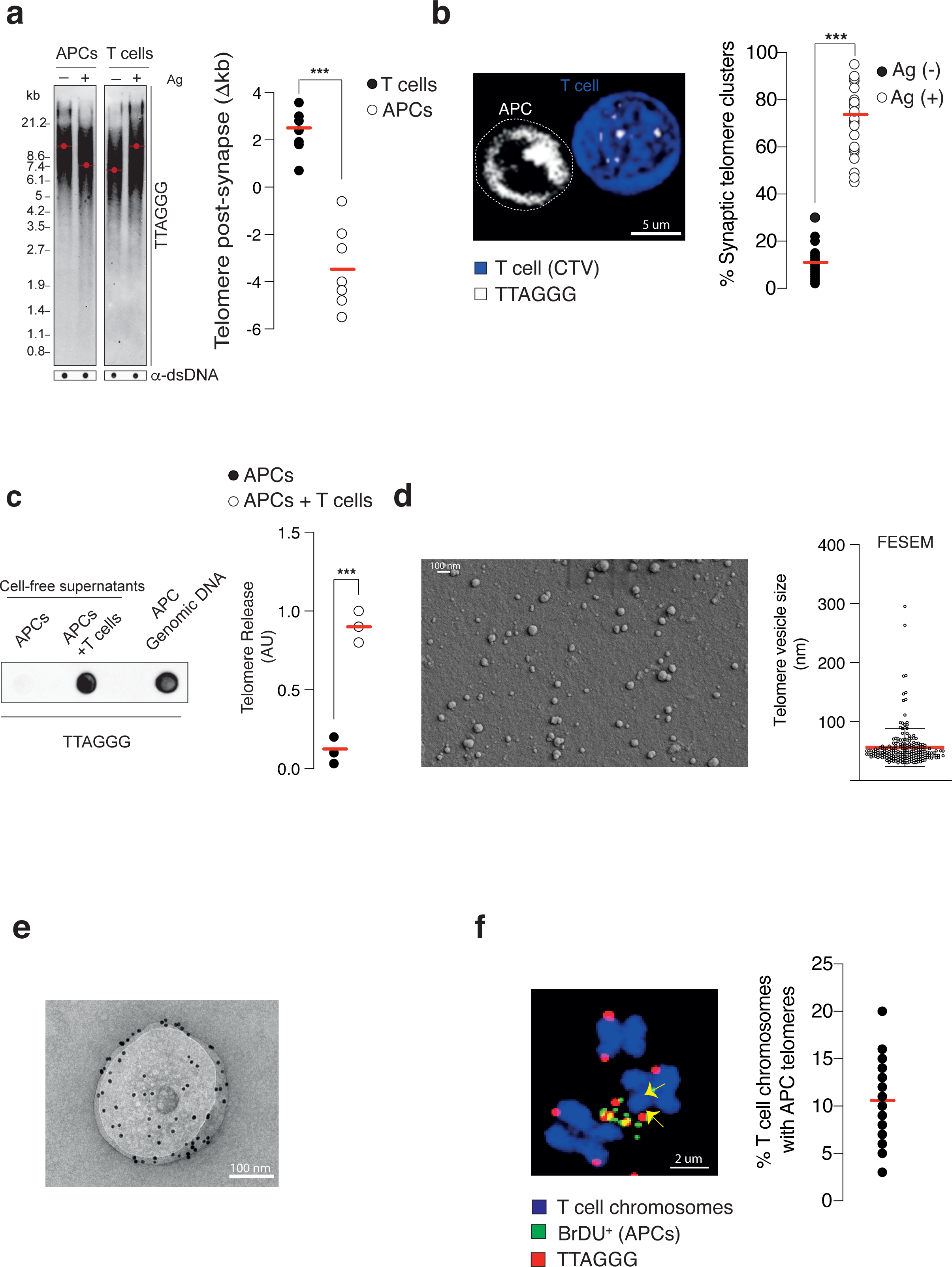
APCs donate telomeres to T cells. (**a**) APCs and T cells were left either unconjugated or conjugated in the presence of 1:50 dilution of an antigen mix (HBV, influenza and CMV lysates), conjugates disrupted, and T cells and APCs isolated by anti-CD3. TRF analysis with TelC probe demonstrated telomere elongation in T cells and reciprocal telomere shortening in APCs 24h later. Red dots indicate mean telomere length. Pooled data from 7 donors are shown (**right**). Dot blots with anti-double strand DNA, loading control. TTAGGG, telomere sequences detected by the TelC probe throughout experiments. (**b**) Telomere clustering in APC-T cell conjugates. APCs were conjugated with Cell Trace Violet (CTV) T cells for 2h then fixed and analysed by IF-FISH with TelC probe. Quantification of telomere clusters in 37 microscopy fields (5-10 conjugates per field) in the presence or in the absence of antigen mix are shown (**right**). Scale bar, 5 μm. (**c**) Telomere donation by APCs. APCs were labelled with BrdU prior to the addition of unlabelled T cells, in the presence of antigen mix. Telomeric DNA donated by APCs was then immunoprecipitated with anti-BrdU from cell-free supernatants and telomeres detected by dot-blot with TelC probes. Input was APC genomic DNA (200 ng). Pooled data from 3 experiments are shown (**right**). Cell-free supernatants are obtained by sequential 300g, 2,000g and 10,000g centrifugation throughout experiments. (**d**) Ultrastructural analysis of telomere vesicles. APCs were live-labelled with the membrane dye PKH67 and TelC telomere specific probes, treated with ionomycin (0.5 μg/mL) for 18h, and telomere vesicles were purified from APC supernatants by fluorescence activated vesicle sorting prior to field emission scanning electron microscopy (FESEM). **See also Extended Data** Fig.2**,3.** Representative FESEM micrograph (**left**) and pooled data depicting the size of 204 purified telomere vesicles at magnification of 100,000X are shown (**right**). Scale bar, 100nm. (**e**) Transmission electron microscopy (TEM) of telomere vesicles. APCs were live labelled with biotin-labelled TelC probe, stimulated with ionomycin, and supernatants subjected to sequential centrifugation on sucrose gradient. Vesicle pellets recovered at 100,000g were then analysed by TEM after fixation, permeabilization and incubation with 10nm streptavidin-gold conjugated antibody. Scale bar, 100nm. (**f**) APCs were labelled with BrdU as above, treated with ionomycin, and the telomere rich cell-free supernatants were transferred to separate T cell cultures that were not labelled with BrdU. T cells were then activated with anti-CD3 plus anti-CD28 (each at 0.5μg/mL) for 48h and metaphase spreads were generated by treatment with colcemid (0.2 μg/mL) for additional 12h, fixed, and analysed by IF-FISH with anti-BrdU and TelC probes. Telomeric DNA donated by APCs was detected at ∼10% T cell chromosomes ends. The pooled data from 300 metaphase spreads are shown (**right**). Scale bar, 2 μm. In (**a, c**), paired Student’s t test. \*\**P* < 0.01. In (**b**), Mann-Whitney test. Error bars indicate S.E.M. throughout.

The differential telomere signal was not due to telomerase reactivation because TERT deficient T cells, in which ∼70% telomerase activity was eliminated by short-RNA interference directed to TERT (the catalytic sub-unit of telomerase), still elongated telomeres upon interactions with APCs (**Extended Data Fig. 1c**). To eliminate all telomerase activity, we generated TERT-KO T cells by clustered regularly interspaced short palindromic repeats (CRISPR) technology. TERT-KO T cells elongated telomeres after exposure to APCs in the absence of any detectable telomerase activity (**Extended Data Fig. 1d**), confirming telomerase-independent telomere elongation.

We next considered whether telomerase-independent telomere replication was due to alternative-lengthening of telomeres (ALT)^7^. However, we found that telomere elongation in both telomerase positive and negative T cells occurred in the absence of DNA synthesis at both strands of telomeric DNA (C- and G-rich strands; **Extended Data Fig. 1e**). Chemical ablation of DNA synthesis in T cells treated with the DNA polymerase inhibitors aphidicolin and thymidine prior to incubation with APCs confirmed that telomere elongation did not require DNA synthesis (**Extended Data Fig. 1f**).

Before synapse formation, APCs had telomeres that were ∼3kb longer than T cells. Because telomere elongation in T cells occurred independently of telomerase or DNA replication, we considered whether APCs would donate telomeres upon synapse with T cells. Using IF-FISH, we discovered that APCs polarised their telomeres into clusters at ∼70% antigen-specific conjugates (**Fig. 1b**), while less than 10% of the conjugates displayed APC telomere clusters in the absence of antigens. No telomere clusters were detected if APCs were not coupled with T cells. Clusters of molecules at the synapse often precede their transfer into the other cell of the conjugate^8^. In fact, using APCs with fluorescently labelled telomeres (**Extended Data Fig. 2**) and 3D-synapse reconstruction, we visualized telomere clusters exiting the nucleus of APCs (**Extended Data Fig. 3a**). Therefore, upon antigen-specific contacts with T cells, APCs organise their telomeres into clusters at the synapse that precedes telomere transfer. APCs also released telomeric DNA to the culture medium upon incubation with T cells (**Fig. 1c**). The secreted telomeres are likely to be encapsulated in lipid vesicles as the telomeric DNA is destroyed by DNase I in the presence, but not in the absence, of a non-ionic detergent (**Extended Data Fig. 3b**).

**Figure 2.**
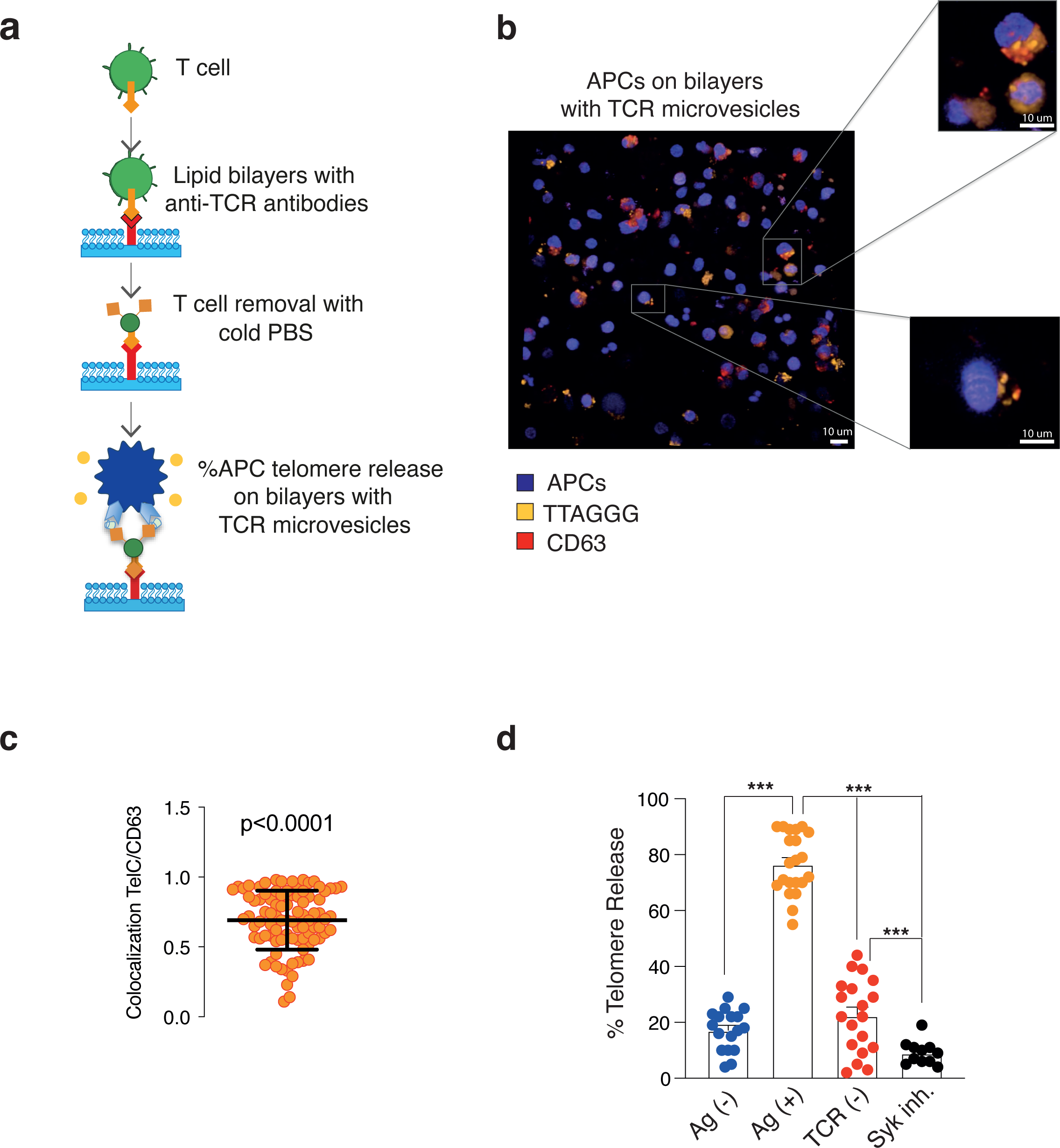
TCRV are sufficient to extract telomere vesicles from APCs. (**a**) Schematic representation of TCRV. T cells were allowed to form synapses on planar bilayers with anti-CD3 and ICAM-1 molecules for 20 min and subsequently removed with cold PBS to retain TCRV on bilayers. APCs were then live-labelled with TelC telomere probes, loaded with antigen mix, and immediately transferred on planar bilayers with TCRV. Telomere release by APCs was quantified 24h later. (**b**) Z-stack 3D reconstruction showing release of CD63^+^ telomere vesicles from APCs activated on bilayers. APCs were live-labelled with TelC telomere probes and anti-CD63 prior to transfer on bilayers. Scale bar, 10 μm. The Z-stack is a screenshot from **Supplementary video 1**. (**c**) Pearson’s co-localization scores between CD63 and TTAGGGs on telomere vesicles released as in (**b**). Each dot is an individual telomere vesicle released by APCs. (**d**) Quantification of telomere release by APCs on bilayers. Data pooled from 4 donors (Ag) or 2 donors (No Ag, No TCR, SYK In). In experiments with SYK inhibitor, APCs were pre-treated with 5 μM SYK inhibitor III for 30 min prior to transfer on bilayers. The fraction of telomeres released by APCs was normalized to the total number of APCs on the bilayer. In (**d**), Kruskal-Wallis with Dunn’s post-correction test. \**P* < 0.05, \*\**P* < 0.01 and \*\*\**P* < 0.001. Error bars indicate S.D. (**c**) or S.E.M. (**d**).

**Figure 3.**
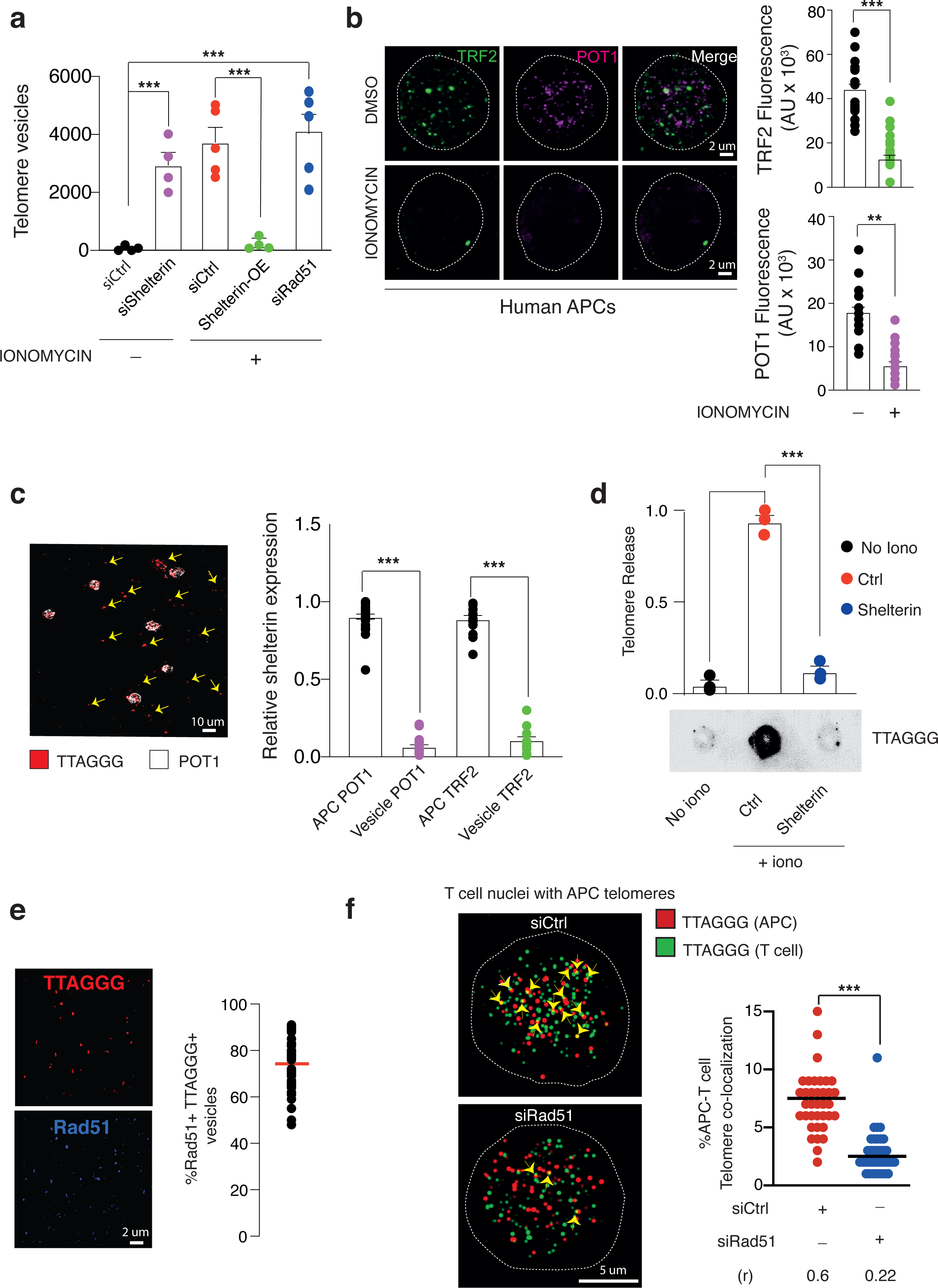
APCs dismantle shelterin to donate telomeres. (**a**) Flow-cytometry quantification of telomere vesicles released by 10^7^ APCs in culture media following the indicated treatments, with or without ionomycin for 18h. (**b**) Shelterin down-regulation in APCs following activation with ionomycin as in (**a**). Representative POT1 and TRF2 staining’s (**left**) and pooled data (n = 2; **rights**) are shown. Each dot is an individual cell. Scale bar, 2 μm. (**c**) APCs donate shelterin-devoid telomeres. APCs were live-labelled with TelC telomere probes, activated with ionomycin, then analysed by IF to shelterin (POT1). Arrows indicate ‘shelterin-devoid’ telomeres released by APCs. Scale bar, 10 μm. Representative microscopy field (**left**) and pooled data from 30 fields (**right**) are shown. (**d**) APCs were transduced with mock or shelterin expressing (TRF2 + POT1) vectors, and labelled with BrdU. APCs were then activated 18h by ionomycin and telomere vesicles released by APCs were immunoprecipitated with anti-BrdU from cell-free supernatants. Representative blots (**bottom**) and data form three experiments (**top**) are shown. **See also Extended Data** Fig.9. (**e**) Telomere vesicles were analysed by IF to Rad51. Individual staining’s (**left**) and quantification from 42 microscopy fields is shown (**right**). Scale bars, 2 μm. (**f**) APCs were transfected with siCtrl or siRad51 for 36h, live-labelled with Cy3-PNA telomere probes then activated by ionomycin for 18h. The cell-free supernatants containing fluorescent APC-telomere vesicles were then transferred into T cells with FITC-PNA telomere labelling. There was no contact between APCs and T cells. APC-T cell telomere co-localization (Cy3 + FITC = yellow) in recipient T cell nuclei was quantified by confocal imaging 24h later. Representative images (**left**) and pooled data from 6 experiments (**right**) are shown. Arrowheads indicate APC-T cell telomere co-localization. Scale bar, 5 μm. In (**a**, **b, c, d, f**), Mann-Whitney test. *p<0.05 \*\*\**P* < 0.001. Error bars indicate S.E.M. throughout.

To further test telomere vesicle hypothesis, we live-labelled APCs using the membrane lipid dye PKH67 and the TelC telomere specific probe, stimulated telomere vesicle release from APCs with ionomycin (for reasons explained below), followed by fluorescence activated vesicle analysis of APC supernatants. Initial attempts were complicated by presence of vesicle clusters that falsely suggested secretion of large extracellular vesicles by APCs. Using a side scatter threshold and calibration beads (**Extended Data Fig. 3c**)^9, 10^, we successfully excluded aggregates and visualized PKH67^+^ TelC^+^ telomere vesicles within ∼10% of the total APC single particle vesicle fraction (**Extended Data Fig. 3d**). qPCR, with telomere primers that ensure no non-specific amplification, confirmed that purified telomere vesicles (Tel^+^) contained telomeric DNA. By contrast, telomere depleted vesicles did not contain telomeric DNA (Tel^-^). Super-resolution microscopy confirmed that PKH67-labelled membrane lipids and TelC-labelled telomeric DNA co-localized in purified Tel^+^ vesicles released by APCs (**Extended Data Fig. 3e**). Field Emission Scanning Electron Microscopy (FESEM) revealed that most Tel^+^ vesicles had a size distribution of 56 +/- 21 nm, characteristic of exosomes (**Fig. 1d**). Using transmission electron microscopy (TEM) coupled to immunogold based telomere detection, we detected telomeric DNA within vesicles (**Fig. 1e**), and confirmed that most telomere vesicles were exosome-sized, with a minor telomere vesicle subset (∼5%) between 100-300 nm characteristic of plasma membrane derived microvesicles (**Extended Data Fig. 3f**). The amount of telomeric DNA detected in the exosome-sized vesicles of 30-100 nm was ∼2 copies per vesicles, while larger vesicles in the 100-300 nm size range contained ∼45 copies per vesicle, which is proportional to volume. Dot blot further showed that telomeric DNA is mostly present in the APC exosome fraction as double-strand (ds) DNA filament (**Extended Data Fig. 4a**). Telomeric restriction fragment analysis, in the presence or in the absence of restriction enzymes to digest non-telomeric DNA, demonstrated that the telomere vesicles have an average telomere length of ∼5kb, not bound to other forms of DNA (**Extended Data Fig. 4b**). Taken together, telomere vesicles are a new class of extracellular vesicles dedicated to the transfer telomeric DNA.

**Figure 4.**
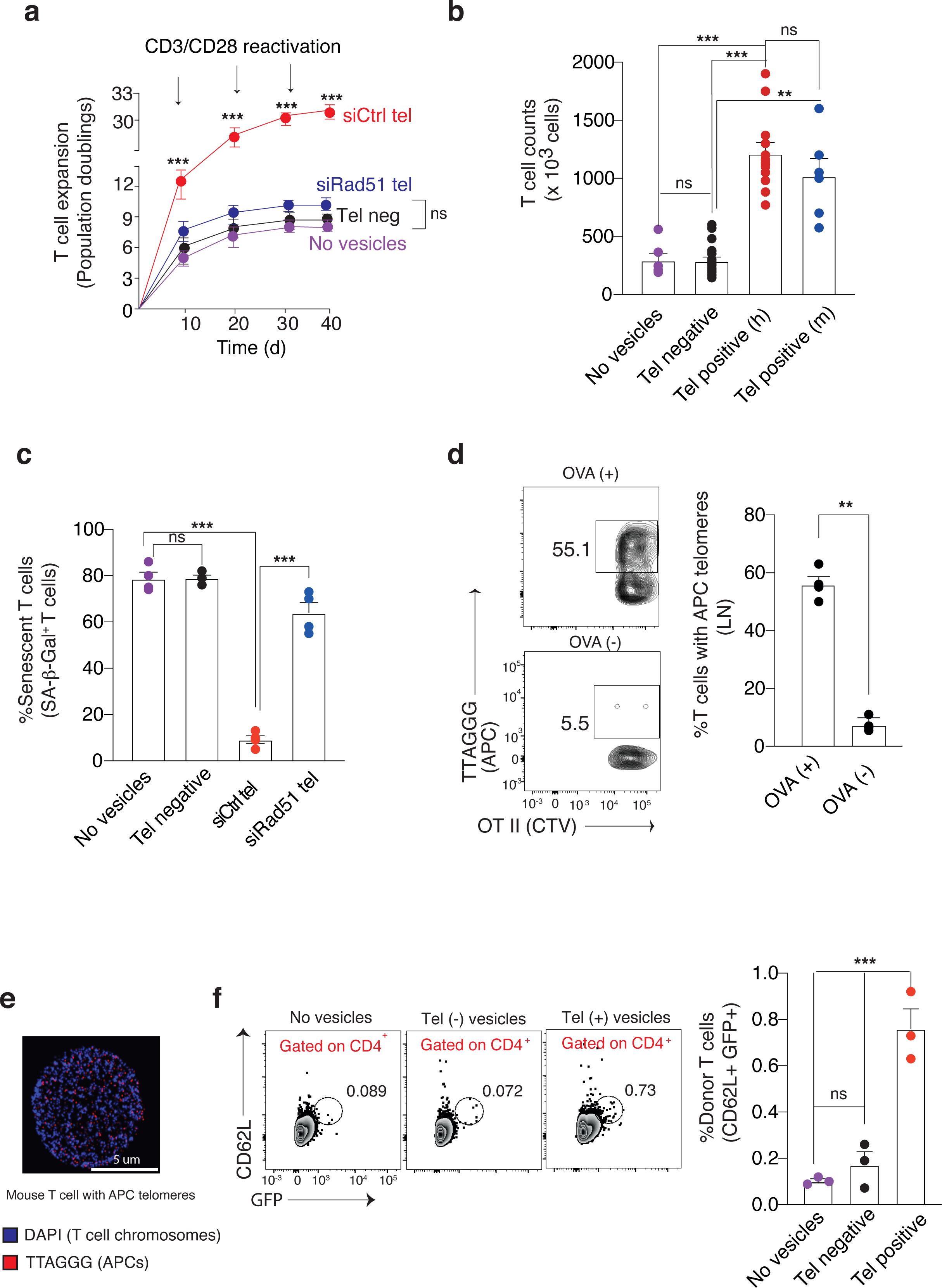
APC telomeres extend T cell lifespan. (**a**) T cells were activated with anti-CD3 plus anti-CD28 every 10 days in the absence or presence of 1,000 siCtrl or siRad51 telomere vesicles (Tel^+^) purified from APCs by FACS. Control T cells were cultured with 1,000 telomere depleted vesicles (Tel^-^) or left without any vesicles (no vesicles) throughout. Population doublings, PD. The experiments were stopped after 39 days at which point the control T cells stopped dividing (∼10 PD). (**b**) Expansion of human T cells by heterologous telomere vesicles. Control T cells were cultured 10 days in the presence of 1,000 telomere vesicles (Tel^+^) or telomere depleted vesicles (Tel^-^) purified by FACS as in (**a**) and derived from either mismatched human (h) or mouse (h) APCs, as indicated. T cells cultured without any vesicles are also shown. Each dot is an independent T cell culture. (**c**) Senescence-associated beta-galactosidase activity was measured in T cells activated as in (**a**) for 10 days. Four donors. (**d**) OVA-pulsed APCs with Cy-3 TelC labelled-telomeres were injected into the right footpad of recipients followed by adoptive-transfers of OT-II cells (identified by Cell Trace Violet dye). Animals were culled 18h after the addition of T cells and popliteal lymph nodes collected. The frequencies of donor OT-II cells with APC-derived telomeres were calculated by flow-cytometry. In control experiments, APCs with fluorescently-labelled telomeres were not loaded with OVA antigens. Representative plot (**left**) and results from 3 experiments are shown (**right**). (**e**) Presence of APC-derived telomeres into the nuclei of mouse T cells (metaphase spreads) following *in vivo* adoptive transfer experiments performed as in (**d**). Scale bar, 5 μm. (**f**) GFP^+^ (donor) OT-II cells were pre-treated with or without 250 telomere vesicles derived from mouse APCs activated with ionomycin, injected into recipients, and splenocytes were analysed by flow cytometry with anti-CD4 and anti-CD62L 8 weeks post-transfer (14 days after re-challenge with OVA antigen, **left**). Controls were injected with either 250 telomere depleted vesicles (Tel-) or saline solution (no vesicles). Pooled data from 3 mice per group are shown (**right**). In (**a, b, c, f**), ANOVA with Bonferroni post-test correction; in (**d**), paired student T test. \*\*\**P* < Error bars indicate S.E.M. throughout.

We next studied the protein composition of telomere vesicles. Using immunoblotting, we detected exosome markers (CD63, TSG101), microvesicle proteins (C1q), histocompatibility antigens (MHC II) and telomere specific proteins (TZAP) in the APC vesicle fraction subjected to ultra-centrifugation (**Extended Data Fig. 4c**). By contrast, the mitochondrial marker cytochrome c was firmly retained within the APC cellular fraction, which confirmed lack of cellular contamination in the vesicle preparations. Enzyme-linked immunosorbent assays (ELISA) revealed that exosome markers and histocompatibility antigens are found in both Tel^+^ and Tel^-^ vesicles (**Extended Data Fig. 4d**). By contrast, the telomere-trimming factor TZAP was exclusively found in the Tel^+^ vesicles, suggesting a possible role for TZAP in telomere cleavage from APC chromosomes. Indeed, silencing TZAP by siRNA inhibited telomere vesicle release from APCs (**Extended Data Fig. 5a, b**). Furthermore, using APCs with fluorescently labelled TZAP, we confirmed that TZAP was donated by APCs at immune synapse with T cells and co-localized with the transferred telomeres (**Extended Data Fig. 5c, d**). Thus, APCs donate telomere vesicles via TZAP.

We investigated whether T cell chromosomes incorporated APC telomeres. To exclude APC-contamination, we indelibly labelled APC specific DNA with the thymidine analogue Bromodeoxyuridine (BrdU), and incubated purified BrdU^+^ telomere vesicles with T cells. T cell metaphase spreads imaged by anti-BrdU and IF-FISH demonstrated that ∼10% of T cell chromosome ends contained telomeric DNA donated by APCs (**Fig. 1f**). No BrdU signal was present in metaphase spreads from T cells not exposed to APC vesicles. We repeated experiments using live APCs with fluorescently labelled telomeres. As expected, ∼10% T-cell chromosome ends with fluorescent telomeres of APC-origin were observed (**Extended Data Fig. 6a**). T cell chromosomes with APC-derived telomeres were destroyed upon application of T7 endonuclease (an enzyme that specifically cleaves sites of DNA integration^11^ directly to the metaphase spreads (**Extended Data Fig. 6b**), which indicated integration of APC-telomeres with T cell chromosomes. Using T cell plasma membrane purification, we confirmed that APC-telomeres were not immobilized on the synapse (**Extended Data Fig. 6c**). Instead, APC-telomeres were detected in the nuclear fractions of the T cells. Moreover, APC-telomeres co-immunoprecipitated with the telomere protein POT1 in the T cells (**Extended Data Fig. 6d**).

To directly investigate fusion between T cell and APC telomeres, we transferred purified telomere vesicles from APCs into T cells followed by qPCR. This confirmed telomere elongation in T cells that received telomere vesicles (**Extended Data Fig. 7a**). No telomere elongation could be detected after transferring telomere devoid vesicles. We also assessed whether MHC II molecules present in the telomere vesicles were required for telomere entry into T cells. However, no difference in APC-T cell telomere fusion in the presence of blocking MHC II antibodies could be detected.

Next, we studied which T cell synapse component triggered telomere transfer. T cells release synaptic TCR-enriched extracellular vesicles (TCRV) that trigger antigen-specific calcium signals upon binding MHC II molecules on APCs^12^. Using APC-surrogate synapse planar bilayers constructed with antibodies to CD3 and the integrin-molecule ICAM-1 (**Fig. 2a**), we found that TCRV were sufficient to induce antigen-specific release of CD63^+^ telomere vesicles from APCs, even in the absence of T cells (**Fig. 2b, c** and **Supplementary video 1**). Using brightfield illumination, we confirmed that CD63^+^ telomere vesicles were released on the bilayer (**Extended Data Fig. 7b**). By contrast, telomere vesicle release was suppressed on bilayers prepared without anti-CD3, where TCRV were not released, or if APCs were not loaded with antigens, where MHC II molecules on APCs cannot bind to the subset of antigen-specific TCR in the TCRV (**Fig. 2d**). Therefore, although MHC on telomere vesicles is not required for telomere delivery to T cells, telomere vesicle release required pMHC signalling in the APCs in response to TCRV.

TCRV activate antigen-specific calcium signals in APCs^12^. We therefore investigated whether calcium signalling in APCs is needed for telomere transfer. Telomere vesicle release was abrogated by a Syk inhibitor, which blocks calcium signalling in APCs (**Fig. 2d**). By contrast, as mentioned above, we circumvented the need for antigen recognition by pharmacologically activating APCs with the calcium ionophore ionomycin. While resting APCs spontaneously release little or no telomere vesicles, we found that APCs activated by ionomycin release ∼5,000 telomere vesicles per 10^7^ cells without any evidence of cell death, blebbing, nor need to use T cells or TCRV for APC stimulation (**Fig. 3a** and **Extended Data Fig. 8a-c**). Immunoblotting and TRF following TZAP immunoprecipitation showed that ionomycin increases both TZAP expression and telomere-trimming activity in APCs (**Extended Data Fig. 8d, e**). Therefore, an intracellular rise in calcium stimulates TZAP and consequent telomere cleavage in APCs.

Shelterin is the telomere-binding complex that protects telomeres from the DNA damage response. Consistent with TZAP cleavage at telomeres with reduced concentration of shelterin^13^, APCs activated overnight in the presence of ionomycin down-regulated shelterin expression by 50-80% (**Fig. 3b**), releasing telomeres devoid of both telomere-binding and shelterin-assembling factors, POT1 and TRF2^14, 15^ (**Fig. 3c**). Similar results were obtained upon conjugation of CMV-loaded APCs with T cells to form antigen-specific synapses (**Extended Data Fig. 9a**). Treating APCs with the proteasome inhibitor MG-132 prevented shelterin down-regulation, suggesting that APCs degrade shelterin through the proteasome after forming the synapse, consistent with regulation of shelterin by calcium-dependent proteasomal degradation^16^. Artificially-restoring shelterin binding to telomeres by lentiviral vectors in ionomycin activated APCs (**Extended Data Fig. 9b**), demonstrated that shelterin overexpression reduces telomere vesicle release from ∼5000 to 100 in APCs treated with ionomycin (**Fig. 3a, d**). Consistently, ablation of shelterin by siRNAs to POT1 and TRF2 in APCs (**Extended Data Fig. 9c**) induce the release of ∼5000 telomere vesicles even in resting state (no ionomycin activation; compare **Fig. 3a** and **Extended Data Fig. 9d**). Thus, shelterin protects telomeres not only from DNA-damage but also from telomere transfer. Upon shelterin degradation, TZAP, the telomere-specific trimming factor which binds to telomeres when shelterin levels drop^13^, leads to telomere excision and concomitant telomere vesicle release.

Telomere vesicles retained Rad51, the homologous recombination factor involved in telomere elongation^17^ (**Fig. 3e**). Rad51 and other DNA damage factors (γH2Ax and BRCA2) were also detected in APC vesicles analysed by immunoblotting, whereas BRCA1 was mostly retained within the APC cellular fraction (**Extended Data Fig. 10a**). To study the function of Rad51 in the telomere vesicles, we transfected APCs with non-targeting siCtrl or siRad51 RNAs (**Extended Data Fig. 10b**), labelled telomeres with TelC Cy3 telomere probes, and activated APCs with ionomycin to generate fluorescent telomere vesicles devoid of Rad51. Although electron microscopy inspection of siRad51 telomere vesicles did not reveal structural alterations (**Extended Data Fig. 10c**), elimination of Rad51 from the telomere vesicles also resulted in depletion of BRCA2 (**Extended Data Fig. 10d**), the homologous recombination factor that binds to and assists Rad51 in telomere replication^18^. By contrast, exosome membrane proteins, such as CD63 and MHC class II molecules, or the telomere trimming factor TZAP, were not affected by Rad51 depletion. This identifies specific defects in recombinogenic molecules in telomere vesicles depleted of Rad51. Overall, telomere vesicle production remained unaffected by Rad51 depletion (**Fig. 3a**).

We then transferred siCtrl or siRad51 telomere vesicles to T cells that had been labelled with FITC PNA probes to distinguish T cell from APC telomeres (**Fig. 3f**). APC telomeres from siCtrl vesicles showed ∼10% co-localisation with telomeres of recipient T cells, while APC telomeres from siRad51 vesicles were less than 3% co-localised with T cell telomeres. APC telomeres were present within the interphase nuclei of T cells (prepared from resting T cells), demonstrating that T cell activation was not required for APC-T cell telomere co-localization. To exclude random co-localization, we re-analysed telomere images using the automated Costes algorithm coupled to Pearson’s co-localization test^19^. These automated analyses demonstrated that random telomere images generated by the algorithm never had higher APC-T cell telomere co-localization than the original images. The Pearson’s co-localization score for siCtrl telomere vesicles with the T cell telomeres was 0.6 and for siRad51 vesicles was 0.22, which confirmed disrupted co-localization between APC telomere vesicles and T cell telomeres in the absence of Rad51.

Next, we quantified telomere elongation in individual T cell chromosomes. Using metaphase Q-FISH, we confirmed an average lengthening of 3kb in T cell chromosomes that received siCtrl telomere vesicles from APCs (**Extended Data Fig. 10e**). Conversely, transfer of siRad51 telomere vesicles generated shorter T cell chromosomes, despite comparable telomere length between siRad51 and siCtrl telomere vesicles (**Extended Data Fig. 10f**). qPCR confirmed that the fusion between APC and T cell telomeres requires Rad51 present in the telomere vesicles (**Extended Data Fig. 10g**). Rad51 protects single-strand (ss) telomeric DNA, likely telomere overhangs needed for recombination-based telomere elongation^20^, such that siRad51 telomere vesicles possess much shorter ss-telomeric DNA than telomere vesicles derived from control APCs (**Extended Data Fig. 10h**). Therefore, telomere vesicles depleted of Rad51 possess shorter ssDNA filaments that impinge on APC-derived telomere fusion in T cells.

Although T cells activate telomerase to extend their own telomeres^6, 21^, telomerase reactivation is insufficient for preventing telomere shortening, especially in ultra-short telomeres^22, 23^. Using Universal-single telomere length analysis (U-STELA)^24^, we demonstrated that T cells that received telomere vesicles (but not those receiving Tel-vesicles or CRISPR-mediated telomerase enhancement) reduced their burden of ultra-short telomeres (<3kb) from ∼4 to 1 (**Extended Data Fig. 11a, b**). We therefore investigated whether APC telomeres could support T cell expansion *in vitro*. T cells activated every 10 days by anti-CD3 plus anti-CD28^25^ in the presence of autologous telomere vesicles expanded ∼three-fold compared to cells not receiving telomere vesicles (left either untreated or stimulated with Tel-vesicles) throughout a 39-day experiment (**Fig. 4a**). A single transfer of APC telomeres was sufficient to boost proliferative lifespan of T cells, which was more effective than telomerase activation (**Extended Data Fig. 11c**). These proliferation effects were not observed in T cells exposed to telomere vesicles from siRAd51 APCs (**Fig. 4a**), which are unable to lengthen telomeres in T cells (**Extended Data Fig. 10g**). Strikingly, similarly to autologous vesicles, we found that allogenic telomere vesicles purified from mismatched human and mouse APCs could also support T cell expansion (**Fig. 4b**). Multiparametric flow cytometry analysis showed that human T cells treated with either human or mouse telomere vesicles donated by APCs down-regulated the checkpoint inhibitory receptors PD-1 and CTLA4, the death receptor FAS ligand, and the CD45RO memory marker, but up-regulated the CD28 co-stimulator and the naïve T cell associated CD45RA splice variant that are predictive of high replicative potential (**Extended Data Fig. 12**). No change could be observed upon transfer of telomere-depleted vesicles compared to T cells grown without any vesicle. Therefore, telomere vesicles can be transferred from different donors or species to extend T cell lifespan and associated immune-phenotype.

Next, we investigated the expression of senescence-associated markers in T cells activated in the presence or in the absence of telomere vesicles. Among senescence-biomarkers were beta-galactosidase foci (**Fig. 4c**), a pH-dependent lysosomal activity that generates simple sugars in senescent cells^26, 27^, as well expression of the sestrins (**Extended Data Fig. 13**), a set of ageing-related molecules that suppress replicative lifespan of T cells^28^. The addition of telomere vesicles but not that of telomere depleted vesicles abrogated senescence-associated beta-galactosidase expression compared to T cells grown without any vesicle (**Fig. 4c**). As expected, siRad51 telomere vesicles did not protect T cells from senescence. Thus, senescence in human T cells is prevented upon transfer of telomere vesicles from APCs.

Finally, we sought evidence of intercellular telomere transfer in a whole animal system. Although in-bred mice have much longer telomeres than humans, we found that telomere transfer is conserved in mice, consistent with conserved telomere regulation between human and mouse T cells^29, 30^. Using an antigen-specific (OT II-OVA) system, we transferred APCs with fluorescent telomeres into recipient mice followed by antigen-specific T cells identified by Cell Trace Violet dye. The following day, antigen-specific T cells had acquired fluorescent telomeres from APCs in their popliteal lymph nodes as determined by flow-cytometry analysis (**Fig. 4d**). By contrast, telomere transfer was suppressed if APCs were not pulsed with OVA. Imaging metaphase spreads, we validated *in vivo* uptake of APC-telomeres even into the nuclei of mouse T cells (**Fig. 4e**). Thus, antigen-specific T cells acquire telomeres from APCs *in vivo*.

We next injected equal numbers (2 x 10^4^) of green-fluorescent protein-(GFP) tagged antigen-specific T cells into recipient mice in the presence or absence of APC telomere vesicles. We used both Tel-vesicles and saline solution as control. Animals were then vaccinated with OVA followed by OVA re-challenge 40 days later. Fourteen days after OVA re-challenge, we analysed splenocytes by flow-cytometry with antibodies to the T cell co-receptor CD4 and CD62L (a cell adhesion molecule expressed by long-lived central memory T cells). The frequency of long-lived CD4^+^ CD62L^+^ GFP^+^ T cells (donor cells) was increased by ∼10 fold upon administration of telomere vesicles (**Fig. 4f**). No expansion of T cells could be detected in animals injected with either saline or Tel-vesicles. Therefore, the telomere vesicles boosted T cell lifespan *in vivo*.

In summary, APCs donate telomere vesicles to promote T cell lifespan (**Extended Data Fig. 14**). Thus, while telomerase maintains telomeres throughout T cell divisions ^31^, telomere transfer sets the initial chromosome length during APC-T cell interactions. Why do T cells use telomere transfer in addition to telomerase activation to maintain long telomeres? Our data suggest that telomere transfer is required to limit the burden of ultra-short telomeres in T cells (**Extended Data Fig. 11b**). These ultra-short telomeres are critical for activating the senescence program but may not be eliminated by telomerase^23, 32^. A more substantial telomere elongation than that evoked by telomerase (∼100-200bp) is required to eliminate these ultra-short telomeres: this is ensured by the intercellular telomere transfer program described in this report. It is possible that a loss of telomere transfer capacity rather than a loss of telomerase is responsible for telomere shortening at the basis of ageing. We propose that transfer of telomere vesicles may be implemented in a number of ageing-related pathologies characterized by telomere shortening as well cancer immunotherapy, where it is clear that replicative senescence of T cells present an outstanding medical challenge that cannot be solved simply by activating telomerase^33–35^. Whether telomere transfer may occur universally across the animal kingdom whenever cells interact with each other to modulate their reciprocal lifespan remains to be determined.

## Acknowledgements

We thank Dr. Luca Battistini for support, Prof. Giuseppe Sancesario and Dr. Alfredo Colantoni for reagents and technical advice. This study was supported by the Wellcome Trust (AZR00630) and the Italian Ministry of Health (GR-2018 12365916). M.L.D. was supported by the Wellcome Trust Principal Research Fellowship 100262Z/12/Z. A.N.A. was supported by the Medical Research Council (MR/P00184X/1). M.K. is supported by the NIH (R37AI04477).

## Author contribution

A.L. conceived of, performed, and directed the study, analysed and interpreted the data, provided funding and laboratory infrastructures, formed collaborations, and wrote the paper; B.V. and C.V. designed and performed experiments and analysed individual data sets; S.V. performed lipid bilayer experiments; C.D. and F.M.E. performed cell isolation; V.C. performed immune-phenotyping, cell-death analysis and provided key samples; O.D. performed initial FISH and collected lymph-nodes; M.S. performed fluorescence activated vesicle sorting,; G.B. provided support and reagents; D.W.G. performed *in vivo* manipulations; A.N.A. provided reagents and initial infrastructures; M.L.D. provided lipid bilayer infrastructure, conceptual framework for synaptic vesicles, feedback and advice. M.K. provided feedback, advice, and supported experimental revisions of the final version of the manuscript, which was read, commented and approved by all authors. B.V., C.V., and S.V. contributed equally to this work.

## Materials and Methods

### Human studies

We collected heparinized peripheral blood samples from 140 healthy donors (aged 20-65, male 55% and female 45%). All human specimens were obtained with the approval of the Ethical Committee of Royal Free and University College Medical School, the University of Oxford or the Fondazione Santa Lucia Ethical Committee (CE/prog. 754) with voluntary informed consent, in accordance with the Declaration of Helsinki.

### Cell isolation

Primary human conventional CD27^+^ CD28^+^ CD4^+^ T cells were purified using commercially available kits from Miltenyi. Briefly, peripheral blood mononuclear cells (PBMCs) were isolated by Ficoll followed by negative selection for CD4 (130-096-533). The untouched population was then purified by CD27 microbeads first (130-051-601) yielding CD27^+^ CD28^+^ CD4^+^ T cells. For isolation of mouse T cells from splenocytes, CD4 MicroBeads (L3T4; 130-117-043) were used. For APC purification (CD3^+^-depleted autologous cells), human CD3 MicroBeads (130-050-101) and mouse CD3e Microbead kit (130-094-973) were used to deplete T cells. In BrdU labelling experiments, human APCs were expanded with IL-4 and GMCSF for 48h prior to downstream application to allow BrdU incorporation in APC DNA. In all other cases, APCs were directly used as CD3 negative cells.

### Signalling studies

Telomere vesicles were stained with antibodies to Rad51 (1:100; ab63801, Abcam) and TRF2 (1:300; ab13579, Abcam) and POT1 (1:300; PA-66996, ThermoFisher). Activated T cells were stained with antibodies to sestrin 1 (1:300, ab134091, Abcam). Shelterin expression in APCs was evaluated with antibodies to TRF2 and POT1 as per telomere vesicles.

### Telomerase modulation by CRISPR or short-RNA interference

Human T cells (2-5 x 10^6^) were transfected with 3 μg of TERT KO CRISPR/Cas9 plasmid (h) (sc-400316, Santa Cruz) according to the manufacturer instructions. TERT-KO (GFP^+^) T cells were activated by anti-CD3 plus anti-CD28 and then purified by FACS 96h post-transfection. Control T cells were transfected with 3 μg of control CRISPR/Cas9 Plasmid (sc-418922, Santa Cruz). Knock-out efficiency was confirmed by immunoblots to TERT (1:1000; sc-3093013, Santa Cruz) on purified GFP^+^ T cells. In knock-down experiments, T cells were analysed 48-72h after nucleofection with siRNAs to TERT (10 nM, sc-156050, Santa Cruz) or siCtrl (10 nm, sc-37007, Santa Cruz). In experiments with telomerase enhancement, 10^7^ T cells were transduced with 3 μg of CRISPR telomerase activation particles (sc-400316-ACT) and then used 96h post transduction.

### Telomere Immunofluorescence *In situ* hybridization FISH (IF-FISH)

Sections were fixed in ice-cold 3.7% formaldehyde for 15 min, washed in PBS and treated 20 min with -20 °C ethanol. Blocking was performed for 1 h at room temperature in PBS-TT (8% BSA, 0.5% Tween-20 and 0.1% Triton X-100 in PBS). Primary staining’s were performed overnight at 4 °C with antibodies to TRF2 and POT-1 (1:300; Abcam and ThermoFisher,); CD3 (1:1000; BIO-RAD MCA463G). Secondary staining’s were performed for 1 h at room temperature in the dark with biotinylated goat anti-rabbit IgG AlexaFluor 405 (1:400; A-31556, Life Technologies) and goat anti-mouse IgG1 AlexaFluor 647 (1:800; Life Technologies A-21240, Life Technologies). Cross-linking was performed 20 min in 4% paraformaldehyde. Sections were dehydrated in graded ethanol, treated with DNase-free RNase (100 ng/mL; Thermofisher) for 60’ then hybridized with 40 pM PNA probe solution (Panagene, TelC Cy3, 14 1224PL-01) in hybridization buffer (1M Tris pH 7.2; magnesium chloride buffer; deionised formamide; blocking buffer, Roche; deionised water). DNA denaturation was performed 10 min at 82 °C. Hybridisation was allowed 2 h at room temperature, followed by subsequent washing in 70% formamide, saline sodium citrate (SSC) buffer and PBS, respectively. DNA was stained for 5 minutes in PBS with 1 ug/ml of DAPI and coverslips mounted with Pro-Long Gold (Invitrogen). Imaging was performed using a Leica SPE2 confocal microscope using LAS X version 3.3.0 software (Leica Microsystems, Wetzlar, Germany) or super-resolution ZEISS LSM 800 Airyscan (Zeiss). Sections were z-stacked and the average telomere integrated fluorescence per cell nucleus was quantified by ImageJ software.

### Antigen-specific conjugates

Human APCs were pre-pulsed for 4h at 37 °C with 1:50 antigen mix (HBV, influenza and CMV lysates; bulk telomere measurements) or 1:50 cytomegalovirus (CMV)-lysates (Zeptometrix Corporation; IF-FISH studies). APC-T cell conjugates were then formed by adding autologous CD27^+^ CD28^+^ CD3^+^ CD4^+^ T cells in a 3:1 ratio, spun at 300 rpm for 5’, then incubated at 37 °C as indicated.

### Isolation of extracellular vesicles (EVs) by ultracentrifugation

Vesicles were isolated from APC supernatants following sequential centrifugation (300g 10 min; 2,000g 10 min; 10,000g for 30 min). The 300g centrifugation is used to remove cells; the 2,000g centrifugation removes cell debris; and the 10,000g removes apoptotic bodies and other non-EV structures. Where indicated, the 10,000g centrifuged APC supernatants were further passed through 0.3 μM filters to remove vesicle clusters, and EVs pelleted at 100,000g for 90min to derive exosome-enriched preparations.

### Vesicle Immunoblotting

APCs were activated with ionomycin for 18h prior to being removed by centrifugation at 300g 10min. The APC supernatants were then subjected to sequential centrifugation at 2,000g 10 min; 10,000g for 30 min and, following 0.3 μM filtration, at final 100,000g for 90min. The pellets obtained after each sequential centrifugation step were resuspended in PBS and used for vesicle immunoblotting with the primary antibodies used in ELISA applications (see below) plus p-H2AX S139 (Abcam, ab26350) and BRCA1 (GeneTex, GTX70113). Input was 20% APC whole cell lysates. Fractionation purity was confirmed with antibodies to cytochrome C (BD pharmigen, 556432). Antibodies were used at 1:1000 dilution.

### Generation of APCs with fluorescent telomeres

APCs were adhered overnight on glass coverslips in complete RPMI-1640 medium (supplemented with 10% heat-inactivated FCS, 100 U/ml penicillin, 100 mg/ml streptomycin, 50 μg/ml gentamicin, and 2 mM L-glutamine). Cy-3 TelC-PNA probes were dissolved 1:50 in telomere loading buffer (80 mM KCl, 10 mM K2PO4, 4 mM NaCl, pH 7.2) and gently introduced on adherent live-APCs by rolling pH 7.2. alkali-washed glass-beads (G8772; Sigma Aldrich) for 5 min. Alkali bead washing was performed overnight in 1M NaOH. TelC-Cy3-probe labelled APCs were washed in PBS twice after rolling, and cultured in complete RPMI. Alternatively, APCs were transduced with lentiviral vector encoding fluorescently-labelled telomere-specific protein TZAP (CFP-TZAP; GeneCopeia EX-Z7575-Lv109) for 48h, conjugated with T cells for additional 24h and telomere transfer visualised by IF-FISH.

### Fluorescence activated vesicle sorting

Telomere vesicles were extracted from APCs live-labelled with Cy3 TelC telomere probes and the membrane lipid dye PKH67 (Sigma Aldrich). After labelling, APCs were activated with ionomycin as above described, and the culture media were collected 18h later. After sequential centrifugation (300g, 2000g and 10000g), the APC supernatants were analysed by fluorescence activated vesicle sorting. Fluorescent telomere vesicles (PKH67+ TTAGGG+ single particles) were depleted by MoFlo Astrios-EQ flow cytometer (Beckman Coulter) equipped with 5 lasers (355, 405, 488, 561 and 640 nm wavelengths). To reduce instrument background noise, the system is also equipped with inline sheath filter with 40 nm pore size. The applied pressure to sheath fluid was 60 psi. Routinely alignments were performed with Ultra Rainbow Fluorescent Particles 3.0 µm (Spherotech, URFP-30-2). The triggering threshold was applied to the side scatter on the 488-SSC channel and the relative applied voltage was determined by using size reference beads (Apogee flow systems, 1493) to exclude vesicle aggregates >300 nm. The Instrument setup to identify microvesicles was obtained by balancing triggering threshold and SSC voltage in order to reduce and maintain over time the background noise at 150-300 events/sec.

### DNA content of telomere vesicles

To confirm presence of telomeric DNA in telomere vesicle preparations derived from APCs, 1,000 telomere vesicles were purified by fluorescence activated vesicle sorting and analysed by qPCR using telomere specific primers. Absence of non-telomeric DNA in telomere vesicle preparations was confirmed using random primers (TermoScientific). T cells nuclear extracts served as positive control for non-telomeric amplification.

### Protein cargo of telomere vesicles

Telomere positive and negative vesicles were purified as above described. Protein cargo was analysed by indirect ELISA after overnight incubation of vesicle extracts (obtained by 1% Triton x-100) onto binding plates with a final volume of 50μl. After overnight incubation, plates were washed 3 times in 0.1% Tween-20 in TBS and incubated overnight at 4C with antibodies to: MHC II (Abcam ab55152), TZAP (Abnova H0003-B01P), CD63 (GeneTex GTX28219), BRCA2 (Abcam ab123491), TSG101 (GeneTex GTX70255), Rad51(Abcam ab63801) (all at 1:100 dilution) Background control plates were incubated with PBS. Antibody specificity was confirmed by staining parallel vesicle extracts with IgG (isotype control Cell Signaling, 3900S). Primary antibodies were then detected using horseradish peroxidase (HRP) conjugated antibodies (anti-mouse or anti-rabbit IgG, Invitrogen) at 1:1,000 dilution in 5% non-fat dry milk 0.1% Tween-20 in TBS, at room temperature for 2h. After washing secondary antibodies 3 times in 0.1% Tween-20 in TBS, 50 μl TMB substrate (TermoScientific) was added in each well to detect signals for 15 minutes followed by 50μl stop solution (TermoScientific). Absorbance of triplicate wells was detected at 450nm with an Elisa reader.

### Telomere restriction fragment (TRF) analysis

APCs and T cells were purified as described above. Primary human T cells (5x10^6^) were conjugated 1:1 with autologous APCs for 24h in the presence of 1:50 antigen mix. The day after, conjugates were broken using a syringe with cold PBS plus 5mM EDTA and T cell and APC fractions purified by anti-CD3. In parallel, additional 5x10^6^ T cells and APCs were left unconjugated for comparison of telomere length before and after forming the synapse. Pre and post synapse T cells and APC fractions were then pelleted and genomic DNA extracted using the PureLink™ Genomic DNA Mini Kit (K182001, ThermoScientific). Telomere length was analysed using the TeloTAGGG™ Telomere Length Assay (12209136001, Sigma-Aldrich). Briefly, genomic DNA (1.5 μg/sample) was digested with HinF I/Rsa I restriction enzymes for 2h at 37 °C. Digested DNA was then subjected to gel electrophoresis (10h 4V/cm) on a 1% Agarose gel. The gel was denaturated in 0.5 M NaOH, 1.5 M NaCl, neutralized in 0.5 M Tris-HCl, 3 M NaCl, pH 7.5 then over-night transferred by capillarity to an Amersham Hybond-N+ nylon membrane (GE). Transferred DNA on membrane was UV cross-linked, finally hybridized for 3h at 42 °C with DIG-labelled telomere probes and bulk telomere length analysed as recommended by the manufacturer (Telo-TAGGG Telomere Length Kit Assay; Roche). For TRF on APC vesicle preparations, 1.5 μg of DNA (purified from pellets recovered after final 100,000g ultra-centrifugation of APC supernatants) were analysed in the presence or in the absence of HinF I/Rsa I restriction enzymes to digest non-telomeric DNA.

### Flow-FISH

T cells were conjugated with autologous APCs and spun 2 min at 300 rpm. Telomere length was then measured by a modified version of a published fluorescence *in situ* hybridization–flow cytometer method over a 48h time course.

For determination of absolute telomere length (kb) by Flow-FISH, we used a previously generated standard curve formed by cryopreserved samples with known telomere length as examined by Southern blot of telomeric restriction factors (TRFs). This standard curve allows conversions of Flow-FISH MFI values into TRF kb by calculating Molecules of Equivalent Soluble Fluorochrome (MESF) units^36, 37^.

### T cell metaphase spreads

Primary human T cells were activated with anti-CD3 plus anti-CD28 antibodies (0.5 μg/mL) and treated with telomere-rich cell free supernatants derived from separate ionomycin-activated APC cultures. T cells were treated overnight with colcemid (0.2 μg/mL; Life technologies) 48 h after activation. T cell metaphase spreads were swollen 5 min in KCl buffer (12.3 mM HEPES, 0.53 mM EGTA, 64.4 mM KCl), fixed 5 min in methanol: glacial acetic (3:1 solution) then dried 60 min onto pre-warmed glass-slides. For metaphase Q-FISH, T cell chromosomes with APC-telomeres were generated as above described and directly quantified by Q-FISH coupled to TFL-software analysis in parallel with internal standards with known telomere length. For endonuclease analysis, enzymatic digestion was performed 30 min with 1 unit T7 endonuclease (New England Biolabs) at 37 °C. Chromosomes were stained with DAPI. Sections were z-stacked and the %T cell chromosomes bearing APC-telomeres before and after the T7 endonuclease treatment was calculated within a single stack. The total number of APC-telomeres remained unaffected.

### APC-T cell telomere fusion

T cells were cultured for 48h with 5,000 telomere vesicles or 5,000 telomere depleted vesicles (both obtained by fluorescence activated vesicle sorting as described below). In some experiments, blocking antibodies to MHC class II (1μg/mL) were added to the vesicle preparations prior to transfer into T cells, as indicated. T cell nuclear extracts were then analysed by qPCR using telomere specific and standard primers (8918, ScienceCell research Laboratories) using standard 6800 Roche Analyzer.

### *In vivo* Telomere Transfer

Six week-old (wild type) C57BL/6J mice and OT II mice were purchased from Charles River. TelC-Cy3-probe labelled mouse APCs were loaded with 3 μM OVA peptide (323-339; GeneScript) and injected into the right footpad of wild type C57BL/6J recipients. The day after, OVA-specific OT II cells (identified by Cell Trace Violet dye staining) were adoptively transferred, popliteal lymph nodes harvested after additional 18 h and APC->T cell telomere transfer examined by flow cytometry. In control experiments, APCs were not loaded with OVA antigens.

### Recall studies

OT II cells were isolated from 6-week old OT II transgenic mice. Cells were activated *in vitro* with plate-bound anti-CD3 antibodies plus anti-CD28 antibodies and transduced with green-fluorescent protein (GFP) mock vectors (MOI = 10) for stable tracking. Ten days after transduction, 2 X 10^4^ GFP^+^ transduced OT II cells were adoptively transferred into wild-type C57BL/6J recipients by intravenous injection in the presence or absence of autologous APC-derived telomeres. Mice were subcutaneously vaccinated with 30 μg OVA antigen in 0.1% LPS alum (Thermofisher) one day after the adoptive transfer of T cells. Forty days later, mice were re-challenged as above; GFP^+^ donor T cells were examined by staining with antibodies to CD4 and CD62L after additional 14 days (eight weeks post-transfer).

### Measurement of Telomerase activity

Telomerase activity was assessed with a TeloTAGGG telomerase ELISA kit according to the manufacturer’s instructions (Roche) and extracts of 2 × 10^3^ viable T cells as described previously^27^.

### Long-term cultures

Primary human T cells were activated by anti−CD3/anti-CD28 antibodies (0.5 μg/mL each) every 10 days in the presence or absence of telomere vesicles or telomere depleted vesicles as control and their population doublings calculated as described previously^12^.

### Immunological analysis

Primary human T cells were activated as above described for 10 days in the presence of telomere vesicles (purified from either human or mouse APCs), telomere depleted vesicles or left without any vesicle. Ten days later, T cells were analysed on a Beckman Coulter flow cytometry with multiparametric settings (10 colors). For the immunophenotype characterization, T cells were stained with anti-CD4 Alexa 700 (clone OKT4, Biolegend, Cat. 317626, 1:100), anti-CD27 PerCP5.5 (clone M-T271, Biolegend, Cat. 356408, 1:100), anti-CD28 APC-Vio770 (clone REA612, Miltenyi Biotec, Cat. 130-116-506, 1:100), anti-CD45RA BV421 (clone HI100, Biolegend, Cat. 3104130, 1:100), anti-CD45RO ECD (clone UCHL1, Beckman Coulter, Cat. M2712U, 1:100), anti-PD1 APC (clone NAT105, Biolegend, Cat. 367406, 1:100), anti-FasL PE-Cy7 (clone NOK-1, Biolegend, Cat. 306418, 1:100), anti-CTLA4 PE (clone L3D10, Biolegend, Cat. 349906, 1:100) and anti-TIM-3 VioBright FITC (clone REA602, Miltenyi Biotec, Cat. 130-109-711, 1:50). Cells were washed in PBS and analysed on a Cytoflex flow cytometer (Beckman Coulter).

### Cell death analysis

APCs were activated with ionomycin as above or conjugated with autologous T cells (3:1) in the presence or absence of antigen mix. Cell death was analysed 18h later by FITC Annexin V/PI Apoptosis Detection Kit with PI (640914; Biolegend).

### Measurement of senescence-associated-β-galactosidase

Human T cells were activated by anti-CD3 plus anti-CD28 either in the presence or in the absence of telomere vesicles from APCs for 10 days. After overnight resting at 37 degrees °C, cells were cytospun 5’ at 500 rpm and senescence-associated beta-glalactosidase activity measured as recommended by the manufacturer (Cell Signaling Technology) on a Zeiss-Axio inverted phase-contrast microscope. Control APCs were activated either in the absence of vesicles or with telomere depleted vesicles.

### APC-T cell Telomere co-localization

APCs were nucleofected with siCtrl (10 nM; sc-37007, Santa Cruz) or siRad51 (10 nM; sc-36361, Santa Cruz), labelled with Cy3-TelC PNA probes and extracted as above described. APC-derived telomeres were then added drop by drop into autologous T cells that had been previously labelled with FITC-TelC PNA probes and cultured onto poly-lysine-coated coverslips. APC-T cell telomere co-localization (Pearson’s co-localization test) into the nuclei of recipient T cells was assessed by ImageJ software. Costes method was used to verify co-localization thresholds against randomly generated images^19^. Co-localization frequencies shown in **Figure 3f** were calculated within a single stack with the following formula:

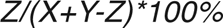

where *Z* is number of co-localization events; *X* is the number of T cell telomeres; and *Y* is the number of transferred telomeres of APC-origin.

### Chromatin Immunoprecipitation

Primary human T cells (10^7^) were activated overnight by anti-CD3 plus anti-CD28 in the presence or absence of Cy3-TelC-labelled APC-derived telomeres (extracted and purified as above described). Cells were cross-linked 10 min with 1% formaldehyde, lysed and their nuclei digested with MNase-based Thermofisher Pierce Agarose Chip kit. Digested chromatin was immunoprecipitated with polyclonal anti-POT1 or control rabbit IgG antibodies at 4C on a rotary shaker overnight followed by incubation with Chip-grade A/G agarose beads at 4C for 3 hours. Presence of APC-derived telomeres in control and POT-1 immunoprecipitated T-cell reactions was quantified as Cy3-absorbance emission of triplicate wells using a micro-plate reader. Presence of APC-derived telomeres was further confirmed by adding DNAse directly to the POT1 IPs for 10 min prior to reading. Background fluorescence was calculated in parallel POT1 immunoprecipitation reactions from activated T cells not treated with APC-derived telomeres.

### Supported planar bilayers

Glass coverslips were cleaned 20 min with Piranha solution, rinsed extensively and dried. Coverslips were plasma cleaned 4 min then assembled into ibidi chambers for bilayer formation. Lipid bilayers containing 12.5% mol NTA (1,2-dioleoyl-*sn*-glycero-3-*N*-5-amino-1-carboxypentyl iminodiacetic acid succinyl) and and 0.004% CapBio (1,2-dioleoyl-*sn*-glycero-3-phosphoethanolamine-*N*-cap biotinyl) in DOPC (1,2-dioleoyl-*sn*-glycero-3-phosphocholine) were incubated 20 min at a total phospholipid concentration of 0.4 mM. After washing in 0.1% HBS/BSA, bilayers were blocked 20 min with 2% BSA/HBS containing 100 µM NiSO4, washed extensively, and loaded 20 min with 4µg/mL streptavidin. Streptavidin-coated bilayers were subsequently coated 20 min with 1.1 μg/ml UCHT1 568 (final density: 30 molecules/ µm^2^) and 0.39 μg/ml ICAM-1 405 (final density: 200 molecules/ mm^2^). Bilayers were washed and immediately used for synapse formation. T cells were activated on bilayers for 20 min, fixed 10 min with 2% PFA then imaged in TIRF mode with Olympus 150x 1.45 NA objective. Primary human T cells were loaded onto bilayers directly *ex vivo* or after over night resting in complete RPMI at 37°C.

### Extraction of APC-telomeres on planar bilayers

Planar bilayers were constructed as described above, and loaded with primary human T cells. After 20 min activation on bilayers, T cells were removed with cold PBS. TCR microvesicles (TCRV) released from the synapse were retained on the bilayer. Next, telomere-labelled APCs were loaded with antigen mix and transferred on the TCRV coated planar bilayers. The % of APCs releasing telomeres on bilayers was quantified 24h later. In control experiments, APCs were not laded with antigen mix; or they were transferred onto bilayers constructed with ICAM-1 molecules alone. The vesicular nature of the released telomeres was further tested by pre-incubating TelC labelled APCs with antibodies to the exosome marker CD63 (clone H5c6; Biolegend Cat. 353015) at 1:50 dilution prior to transfer on bilayers.

### Dot blot analysis of telomere vesicles

Human APCs were pulsed with BrdU (10 μg/mL) and cultured in the presence of IL-4 (50 ng/mL) plus GM-CSF (10 ng/mL) for 48 hours, at which point BrdU had been uniformly up-taken by APCs (data not shown). APCs were then washed with PBS, loaded for 4h with CMV antigens, and allowed to form synapses with BrdU-unlabelled (autologous) T cells. Twenty four hours later, supernatants were collected, purified by sequential centrifugation (as above described), and DNA secreted by APCs was firstly concentrated 60 min at -80 °C by adding Ethanol/sodium acetate, then immunoprecipitated with anti-BrdU in the presence of 1% Triton-X100 at 4°C. The next day, BrdU immunoprecipitates were incubated with Agarose A/G beads (1:100; Santa Cruz Biotechnology) and kept rotating at 4°C for 3h. Beads were subsequently washed twice with ice-cold HNGT buffer (50 mM HEPES, pH 7.5, 150 mM EDTA, 10 mM sodium pyrophosphate, 100 mM sodium orthovanadate, 100 mM sodium fluoride, 10 mg/ml aprotinin, 10 mg/ml leupeptin, and 1 mM phenylmethylsulfonyl fluoride) and once with TE buffer. Beads were then incubated in elution buffer (1% SDS in TE buffer) for 15 min at 65 °C. Telomere vesicles were then eluted by centrifuging at 13,000 rpm for 30 min and dot blotted on Amersham Hybond-N+ nylon membrane (GE). The membrane was denaturated in 0.5 M NaOH, 1.5 M NaCl, neutralized and UV-cross linked for 2 min. Telomere dot blot was performed by 3h hybridization at 42°C with DIG-labelled telomere probe as recommended by the manufacturer (Telo-TAGGG Telomere Length Kit Assay; Roche). Control Input was 200 ng genomic DNA from APCs. Alternatively, telomeric DNA was pelleted from APC vesicles isolated by sequential centrifugation. DNA was sonicated (5 cycles 20 seconds/each) and spotted onto Nylon membranes. Presence of single strand versus double strand telomeric DNA was confirmed by 10mM EDTA plus 100mM NaOH denaturation followed by 10min at 95°C prior to transfer onto Nylon membranes. Pellets were also recovered from all previous centrifugation steps as a control, largely showing absence of telomeric DNA in earlier purification steps.

### Shelterin modulation

Human APCs were cultured with IL-4 (50 ng/mL) plus GM-CSF (10 ng/mL) for 48h, then transduced with POT1 (sc-403275-LAC) and TRF2 (sc-401289-LAC) lentiviral activation particles (both from Santa-Cruz Biotechnology). Control APCs were transduced with control lentiviral activation particles (sc-437282-lac). Four days post-transduction, APCs were activated with ionomycin and analysed by IF-FISH with anti-POT1 and anti-TRF2 antibodies as well telomere dot blot as above described. Conversely, shelterin-deficient APCs were generated by siTRF2 plus siPOT1 nucleofection from 10^7^ human APCs and their telomere vesicle release analysed 72h later as above described.

### Universal-STELA

Briefly, 10 ng of MseI and NdeI digested DNA was incubated with 12-mer and 42-mer panhandles at 65 °C then cooled to 16°C in 49 minutes. T4 DNA ligase was then added to the mixture overnight and kept at 16°C (final volume, 15 μl). The digested DNA was then incubated with 10^-3^ μM terminal adapters (telorettes) in a final volume of 25 μl and incubated overnight at 35°C. Next, a PCR reaction was set using 40 pg (i.e. 5 genome equivalent) of ligated DNA, in the presence of 0.1 μM adapter and teltail primers and using failsafe master mix and failsafe enzyme. The PCR conditions were as follows: 1 cycle at 68°C for 5 min; 1 cycle at 95 °C for 2 min; 26 cycles at 95 °C for 15 sec; 26 cycles at 58 °C for 30 sec; 26 cycles at 72°C for 12 min; 1 cycle at 72 °C for 15 min. The PCR product was then resolved as per TRF analysis on a 0.8% Agarose gel and the load of ultra-short telomeres was calculated as number of telomere bands <3kb/ genome equivalent as described(*19*). U-STELA PCR primers (5’—3’) were as follows: 11+2-mer-panhandle <u>TAC CCG CGT CCG C</u> 42-mer-panhandle TGT AGC GTG AAG ACG ACA GAA AGG GCG TGG TGC GGA CGC GGG

Telorette 1 TGC TCC GTG CAT CTG GCA TCC CCT AAC

Telorette 2 TGC TCC GTG CAT CTG GCA TCT AAC CCT

Telorette 3 TGC TCC GTG CAT CTG GCA TCC CTA ACC

Telorette 4 TGC TCC GTG CAT CTG GCA TCC TAA CCC

Telorette 5 TGC TCC GTG CAT CTG GCA TCA ACC CTA

Telorette 6 TGC TCC GTG CAT CTG GCA TCA CCC TAA

Adapter primer TGT AGC GTG AAG ACG ACA GAA

Teltail primer TGC TCC GTG CAT CTG GCA TC-DIG

### Analysis of telomere single-stranded DNA (ssDNA) by qPCR

Vesicle DNA was purified and analysed by quantitative amplification of ssDNA ^38^. Briefly, DNA was amplified by two PCR rounds (2xPCR). In the first round, 4 ng of DNA was added to a 25 µl of a master mix comprising (final concentration in parentheses): water, 5X Colorless GoTaq® Reaction Buffer (1×), dNTPs (250 µM), telomere primer set (0,25 µl) and GoTaq® G2 DNA Polymerase (0.025 U/µl) and the samples subjected to: 2 cycles; 40°C for 5 min ramp to 72°C, at 2°C per min, and 10 min 72°C. PCR products were purified (QIAquick PCR Purification Kit, 28104, Qiagen) and subjected to a second qPCR round using the Absolute Human Telomere Length Quantification qPCR Assay Kit (8918, Sciencell). Vesicle DNA (4 ng) was also analysed by single round qPCR (1xPCR) using the same kit. Percentage of ssDNA was calculated as:

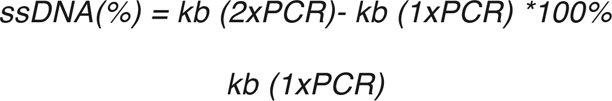

### Field Emission Scanning Electron Microscopy

Samples (APCs or telomere vesicles purified by fluorescence-activated vesicle sorting), were adhered on Ply-L-Lysine-coated round glass coverslips of 1 cm diameter, then fixed with 2.5% glutaraldehyde in 0.1M sodium cacodylate buffer (pH 7.4) at room temperature for 1h. Fixed samples were rinsed 10 minutes with 0.1M sodium cacodylate buffer (3 times), then dehydrated through ethanol gradients (30%, 50%, 70%, 85%, 95%, 100% - 10 mins at room temperature for each ethanol concentration). A 1:1 ethanol: hexamethyldisilazane (HMDS) mix was then added to the samples for 5 min at room temperature followed by a final 5 min step incubation in HMDS. All samples were then left drying overnight. The upper base of each aluminium stub was coated with colloidal silver paint and the coverslips were pasted onto them. Samples mounted on stubs were coated with a gold layer by Q150R S Rotary-Pumped Sputter Coater (Quorum Technologies) and examined by FESEM (Sigma-Zeiss) at an accelerating voltage of 2 kV using secondary electron (SE) detection. A number of regions of interest (ROIs) containing single or multiple APCs for each sample were recorded to confirm absence of glass beads (fig. S2B) or to depict structural alterations (fig. S8C). Micrographs of telomere vesicles were used to perform particle size distribution analysis by measuring the diameter of each adhered telomere vesicle.

### Immunogold analysis of telomere vesicles

Human APCs (10^8^) were live-labelled with TelC-biotinylated PNA probes as above described, then stimulated with ionomycin for 18h. The 10,000g ultra-centrifuged supernatants (1mL) were then loaded over 100 ul of 30% sucrose solution in PBS without mixing the two layers, and ultracentrifuged at 100,000g for 90 minutes at 4 °C using SorvallTM WX 90+ ultracentrifuge (Thermo Fisher Scientific). The supernatants were then discarded and the sucrose layer (∼200 ul) washed with 1 ml ice-cold PBS followed by further ultracentrifugation at 100,000g for 90 minutes. Vesicle pellets were resuspended in 40 μL fixing solution (2% paraformaldehyde, 0.125% glutaraldehyde in 0.1 M sodium phosphate buffer), then fixed for 1h at 4°C on a rocker. 5-μl of vesicle preparations were mounted on Formvar/Carbon 200 Mesh Cu grids (Agar Scientific Ltd), and the grids left to air dry for 10 min. Vesicles were permeabilized with 0.1 % saponin diluted in PBS for 30 min, and washed twice in PBS. Samples were incubated in blocking buffer (1% Bovine Serum Albumin (BSA), 20 mM glycine in PBS) for 45 min, then labelled with Strepavidin-collodial 10 nm gold conjugate (S9059, Sigma Aldrich; 1:10 dilution) in blocking/permeabilizing buffer at 4 °C for 1h on a rocker. The vesicles were washed 6 times in blocking buffer (1:10 in PBS) followed by incubation in 1% (v/v) glutaraldehyde for 5 min and washed in ddH2O. Grids were negatively stained using 1% phosphotungstic acid (Electron Microscopy Sciences) for 1 min and the excess removed using a filter paper. Finally, the grids were air dried for 10 min and examined on a Philips EM 208S (FEI) transmission electron microscope with an accelerating voltage of 80 kV. Images were captured on a MegaView III (Olympus Soft Imaging Solutions). In control experiments, APCs were not labeled with TelC-biotin probes prior to ionomycin-based activation and EV purification that confirmed absence of telomere vesicles with immunogold-labelling (data not shown)

### Exclusion of APC-telomeres in T cell plasma membranes

APC-telomeres were live-labelled with TelC telomere probes, extracted by ionomycin, and transferred to autologous T cells activated by anti-CD3 plus anti-CD28. Twenty-four hours later, T cell plasma membranes were purified with Plasma Membrane Protein Extraction Kit (ab65400; Abcam) and absence APC telomeres assessed by confocal imaging on Leica SP2. Plasma membrane purification was confirmed by CellMask dye (Invitrogen).

### Statistical analysis

GraphPad Prism was used to perform statistical analysis. For pairwise comparisons, a paired Student’s *t*-test was used. For three matched groups, a one-way analysis of variance (ANOVA) for repeated measures with a Bonferroni post-test correction was used. For synapse studies, Mann-Whitney test was used. \**P* < 0.05, \*\**P* < 0.01 and \*\*\**P* < 0.001 and error bars indicate S.E.M. throughout.

### Data availability

The data generated in this study are included in the main paper and its supplementary information files.

---

**Extended Data Figure 1.**
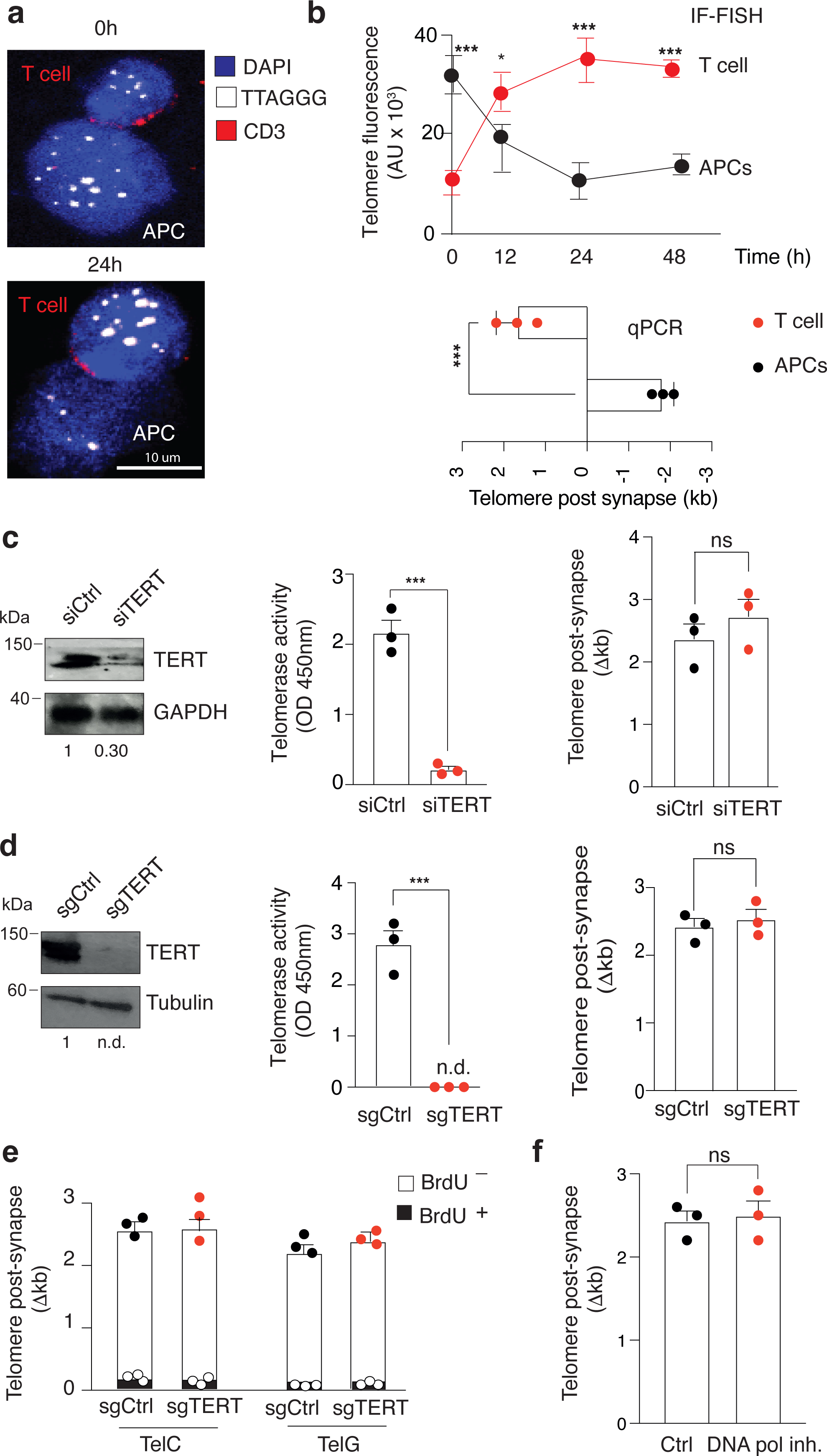
Telomere elongation without DNA synthesis. (**a**) Representative IF-FISH and (**b, top**) pooled data showing telomere elongation in T cells and concomitant telomere shortening in APCs after forming the synapse. T cells were allowed to interact with autologous CMV-loaded APCs for the indicated time points. Conjugates were fixed, and analysed by IF-FISH with TelC telomere probe, DAPI (nuclei) and anti-CD3 (T cells). Three donors. Scale bar, 10 μm. T=0, initial time at which conjugates are observed (20 min). Images were z-stacked and the raw telomere integrated fluorescence signals (AU, arbitrary units) are shown. 102 conjugates were analysed. (**b, bottom**) Analysis of telomere length by qPCR in APCs and T cells after forming conjugates. (**c, d, left and middle**) T cells were transfected with siCtrl or siTERT RNAs (**c**), sgCtrl or sgTERT CRISPR constructs (**d**), activated with anti-CD3 plus anti-CD28, and elimination of telomerase was confirmed by both immunoblot (**left**) and TRAP assay (**middle**). (**c, d, right**) Telomerase positive and negative T cells were exposed to APCs and telomere content was quantified by flow-FISH using TelC telomere probe. Absolute telomere length by Flow-FISH was determined from Mean Fluorescence Intensity (MFI) values using a standard curve formed by cryopreserved samples with known telomere length as determined by TRF. (**e**) Telomerase positive and negative T cells (identified by GFP expression) were pre-incubated with BrdU for 4h, washed to remove any BrdU excess, then stimulated with APCs for 48h. Telomere content was then measured by flow-FISH using either TelC or TelG telomere specific probes coupled to anti-BrdU detection (to monitor telomere elongation *vs* DNA synthesis in T cells). Flow FISH demonstrated that T cells stimulated with APCs elongated telomeres without synthetizing DNA. The T cells remained indeed BrdU^_^ while increasing their telomere content. This suggested that the source of telomeres may be provided directly by the APCs. (**f**) Telomere length by flow-FISH demonstrating telomere elongation in T cells treated with DNA polymerase inhibitors aphidicolin and thymidine (both at 0.5 μg/mL) prior to exposure to APCs for 48h. In (**b, top**) ANOVA with Bonferroni-post test correction, in (**b, bottom**, **c, d, f**) student’s t test. \**P* < 0.05 and \*\*\**P* < 0.001. Error bars indicate S.E.M. throughout.

**Extended Data Figure 2.**
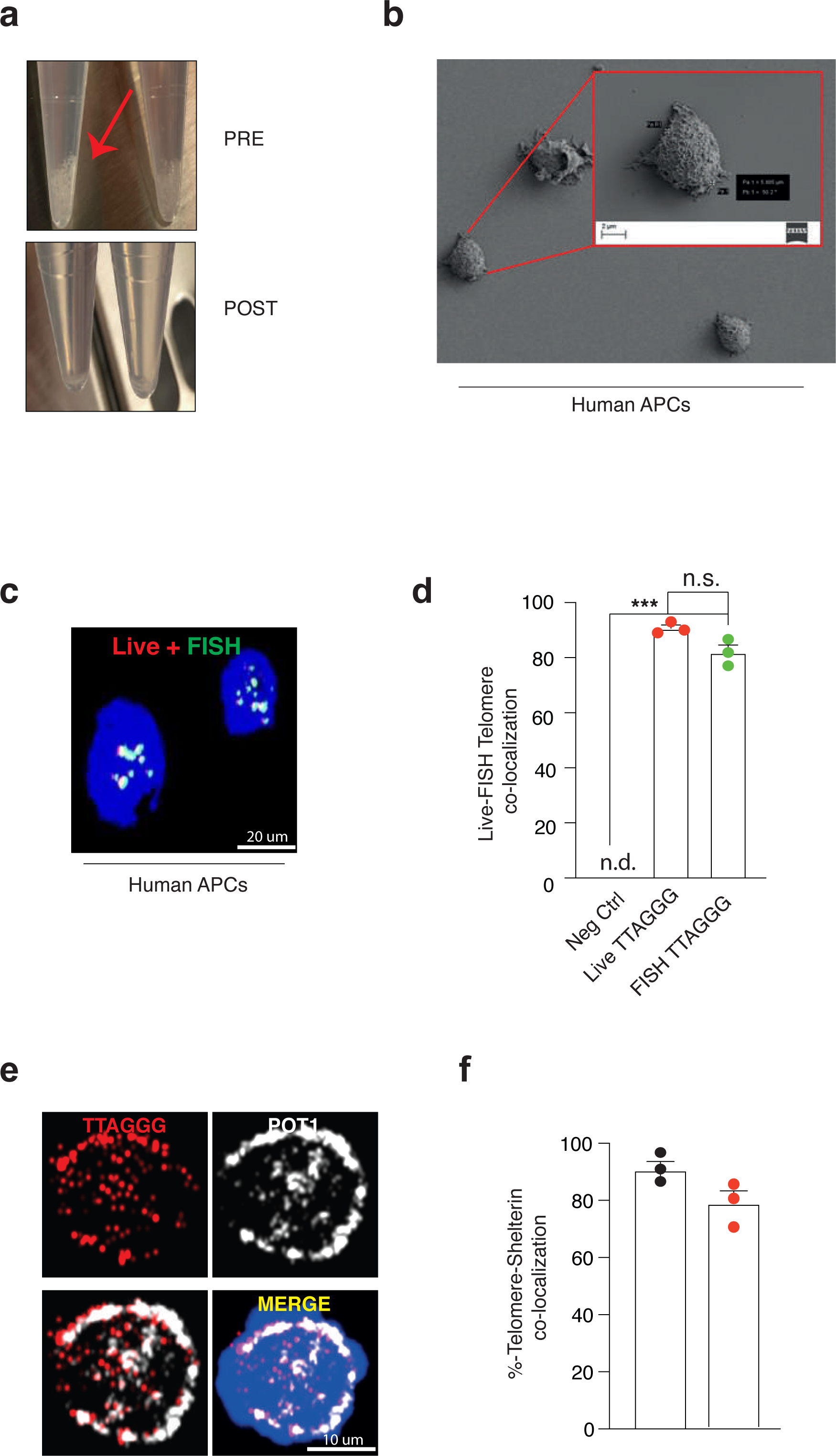
Generation of live APCs with fluorescent telomeres. (**a**) PNA-telomere probes were dissolved 1:50 in telomere loading buffer (80 mM KCl, 10 mM K2PO4, 4 mM NaCl, pH 7.2) and gently introduced on adherent APCs by rolling large glass beads (size: 400-600 nm). The beads are used to produce small holes in the APC plasma membranes that allow telomere labeling while preserving viable APCs (no cell fixative) needed for subsequent synapse studies with T cells. Arrow indicates beads. The beads are completely removed by washing with PBS prior to assays. (**b**) Absence of any residual glass beads from APCs was confirmed by field emission scanning electron microscopy (FESEM). Scale bar, 2 μm. (**c**) Identical telomere detection by IF-FISH (fixed cells) or glass bead-mediated telomere PNA probe delivery (live cells) in APCs. Four experiments. Scale bar, 20 μm. (**d**) Manders co-localization scores of experiments as in (**c**). Negative control, APCs not labelled with telomeres. (**e**) POT1 recruitment to telomeres detected by microbead-mediated telomere PNA probe delivery in human APCs, directly *ex vivo*. Scale bar, 10 μm. (**f**) Manders co-localization scores of experiments as in (**e**). Two experiments. Error bars indicate S.E.M. throughout.

**Extended Data Figure 3.**
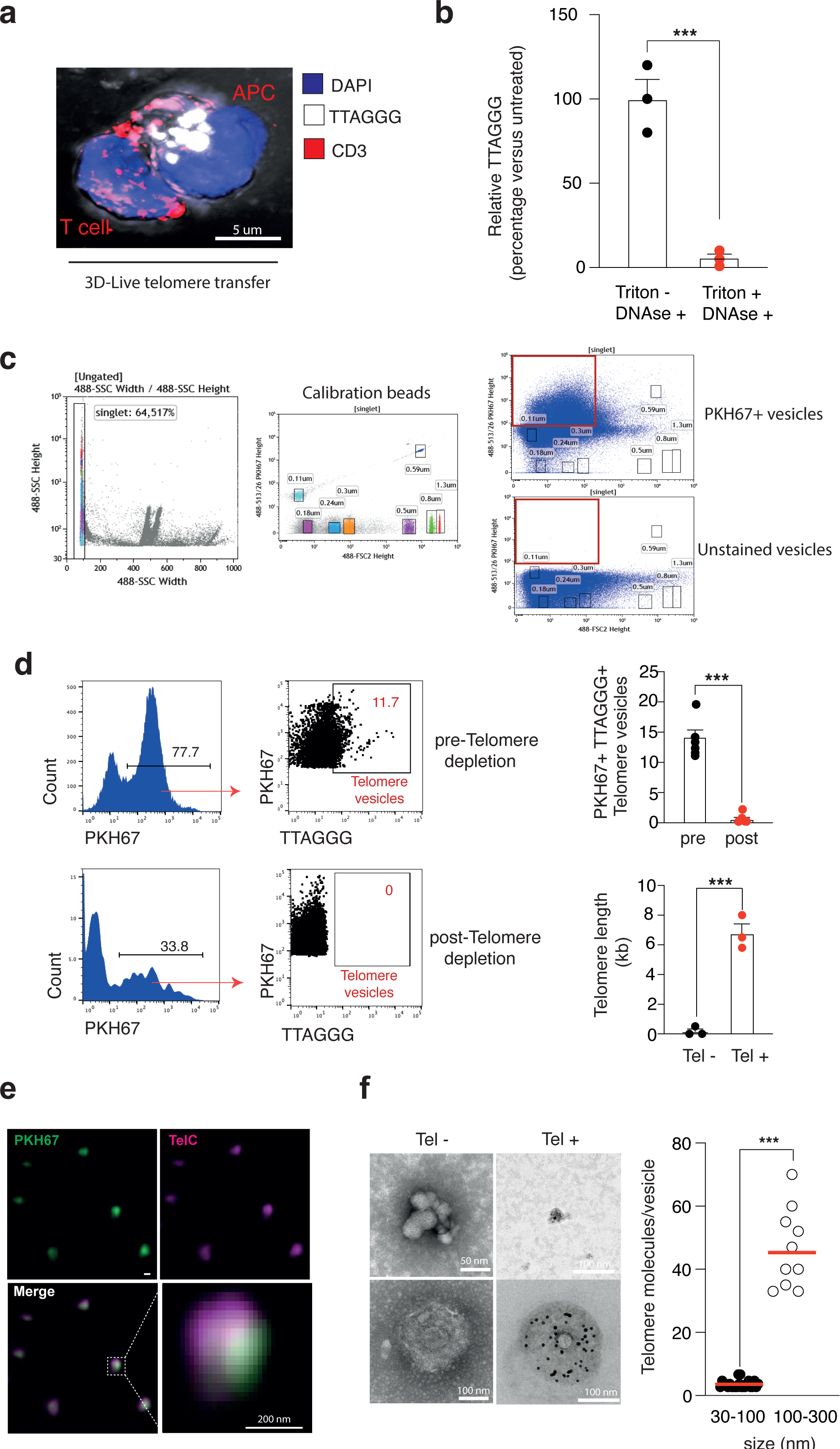
APCs donate telomere in vesicles. (**a**) Z-stack 3D reconstruction showing telomere transfer through the immune synapse. APCs were live labelled with TelC probes then conjugated with telomere-unlabelled T cells for 12 hours. Scale bar, 5 μm. (**b**) Vesicle protection assay. Cell-free supernatants obtained by APCs activated with ionomycin were treated with DNase (10 U, 45’) either in the presence or absence of 1% non-ionic detergent (Triton X-100) then analysed by qPCR. (**c, left and middle**) Side scatter (SSC) threshold and calibration beads used in FACS-based vesicle purifications. (**c, right**). A gating strategy was applied to purify individual vesicles <100 nm up to 300 nm. Larger particles (>300 nm) were instead due to the formation of vesicle aggregates (data not shown) and were therefore excluded from sorting. (**d, left and middle**) Presence of telomere vesicles (TTAGGG^+^ PKH67^+^ single particles) in ∼10% of the total APC single particle vesicle fraction. APCs were live labelled with the membrane lipid dye PKH67 and the TelC telomere probe and activated with ionomycin for 18h. Cell free supernatants were subjected to fluorescence activated vesicle sorting. The representative depletion of telomere vesicles and pooled data from 9 depletion experiments (**top, right**) and the telomere content in purified vesicles (**bottom, right**) are shown. (**e**) Super-resolution microscopy of FACS-purified TTAGGG^+^ PKH67^+^ vesicles (Tel^+^ vesicles). Dashed line indicates vesicle magnification. Scale bar, 200 nm. (**f**) TEM analysis of Tel^-^ *vs* Tel^+^ vesicles performed as in Fig. 1e. Examples for exosome-sized vesicles (**top**) and microvesicles (**bottom**) are shown for both Tel^-^ and Tel^+^. Quantifications of telomeric DNA in exosome-sized vesicles vs microvesicles are shown (**right**). In (**b, d**), student’s t test, in (**f**) Mann-Whitney test \*\*\**P* < 0.001.

**Extended Data Figure 4.**
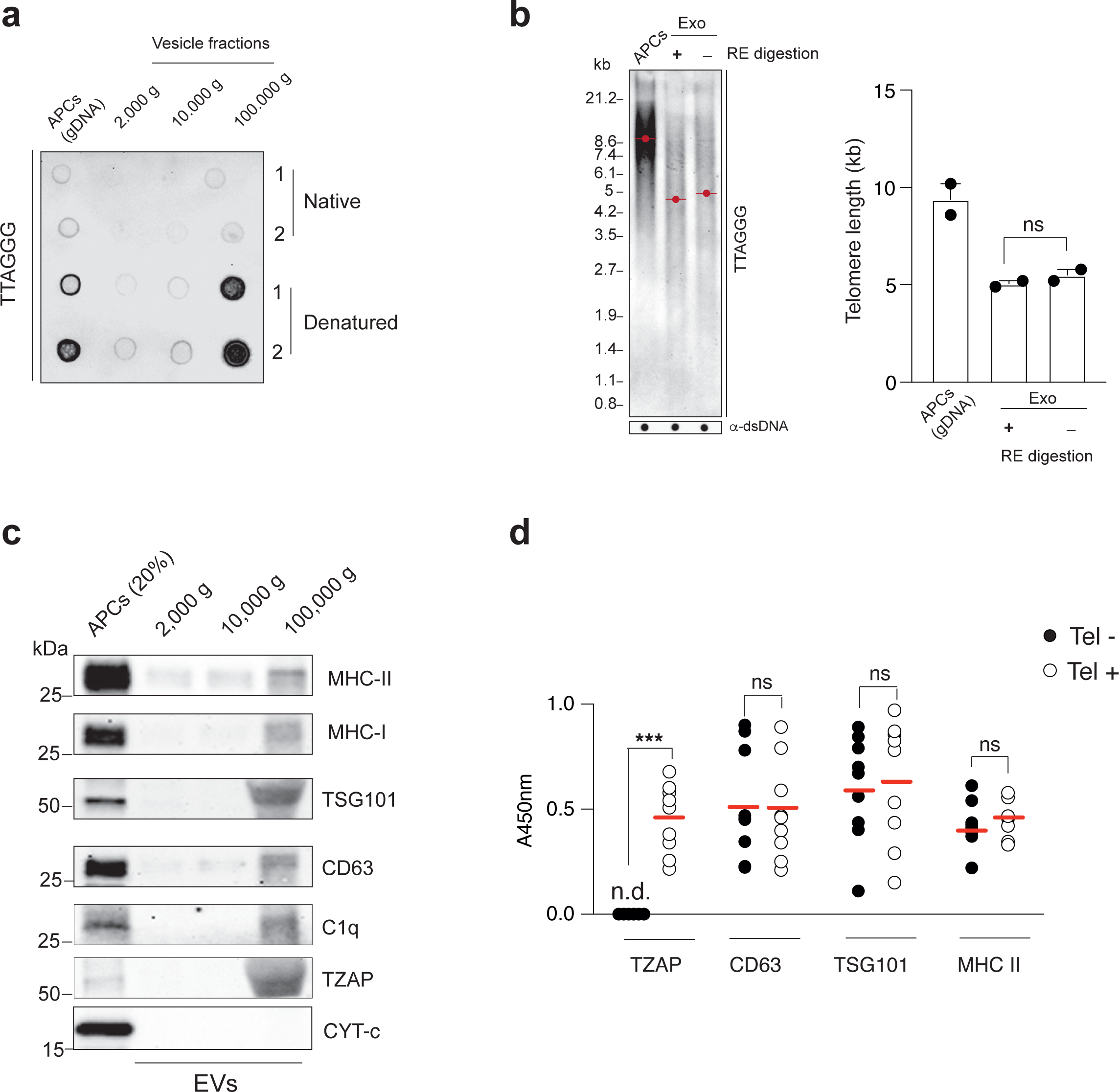
Protein composition of telomere vesicles. (**a**) Dot-blot analysis of telomeric DNA isolated in EV pellets by sequential centrifugation of APC supernatants under native or denaturing conditions. APC genomic DNA (gDNA) served as loading control. (**b**) TRF analysis (**left**) and pooled data (n=2; **right**) of APC telomeric DNA present in the EVs isolated by 100,000 g ultracentrifugation. The addition of restriction enzymes to digest non-telomeric DNA produced no change in telomere length, demonstrating that telomere vesicles released by APCs are largely devoid of non-telomeric DNA. APC gDNA was used as positive control. Dot blots with anti-dsDNA, loading controls. (**c**) Immunoblot analysis of EV pellets derived by sequential centrifugation of APC supernatants after activation with ionomycin for 18h. APC cell lysates (20%) were used as input control. Representative of 3 experiments. (**d**) Protein cargo analysis in FACS purified telomere vesicles (Tel^+^) or telomere depleted vesicles (Tel^-^) by indirect ELISA. The results from nine experiments are shown. In (**d**), student’s t test, \*\*\**P* < 0.001.

**Extended Data Figure 5.**
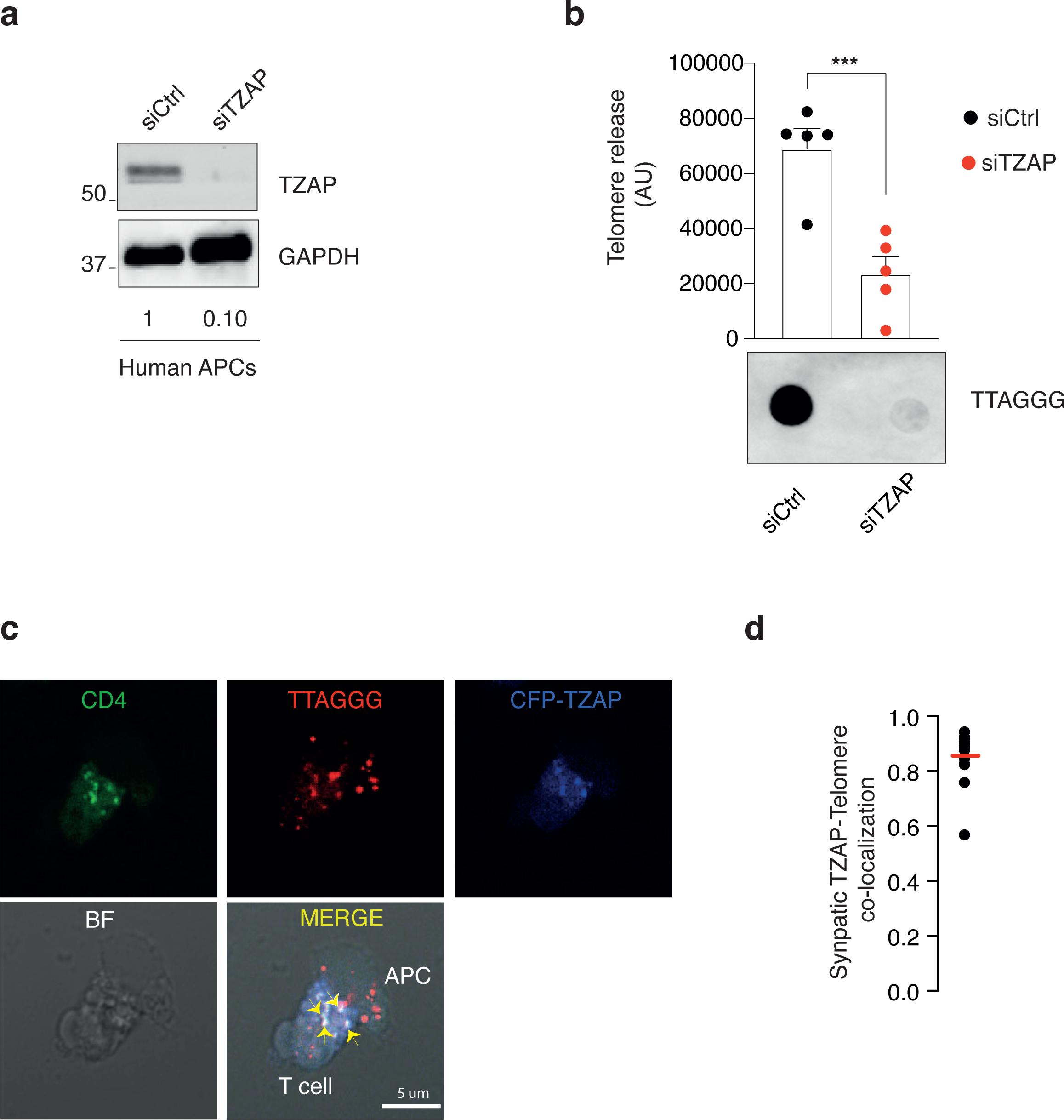
APCs donate telomere vesicles via TZAP. (**a**) Validation of TZAP depletion. Human APCs were transfected with siCtrl or siTZAP for 72h followed by immunoblotting to TZAP. GAPDH served as loading control. The numbers indicate quantification of knock-down efficiency. (**b**) Analysis of telomeric DNA in EV pellets isolated by ultra-centrifugation from APCs transfected as in (**a**) and activated with ionomycin for 18h. Representative results and pooled data from 5 experiments are shown. (**c**) TZAP transfer at the immune synapse. APCs were transduced with lentiviral vectors encoding TZAP tagged to cyan-fluorescent-protein (CFP-TZAP), conjugated 24h with T cells in the presence of antigen mix then analysed by IF-FISH with TelC probe. In APCs transduced with mock vector, no transfer was observed (data not shown). (**d**) Pearson ‘s co-localization score in 15 conjugates as in (**c**). In (**b**), student’s t test, \*\*\**P* < 0.001.

**Extended Data Figure 6.**
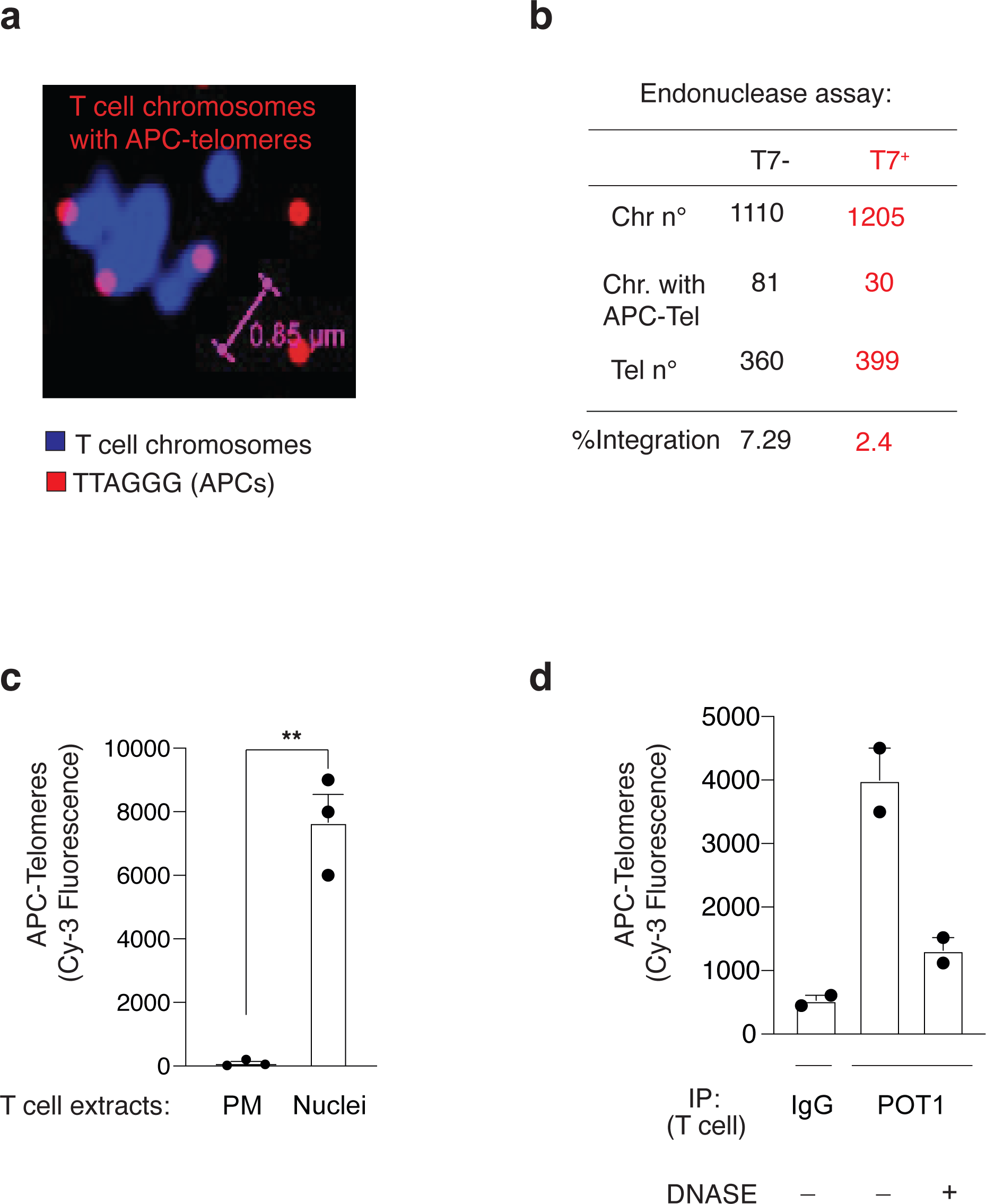
APC telomeres at T cell chromosome ends. (**a**) Representative metaphase spreads showing T cell chromosomes with APC-derived telomeres. T cells were incubated with cell-free supernatants from ionomycin-activated APCs that were live-labelled with TelC probes. Scale bar, 0.85 μm. (**b**) Metaphase spreads generated as in (**a**) were treated with 1unit T7 endonuclease for 30 min at 37 °C. The number of T cell chromosomes with APC telomeres before (black values) and after (red values) T7 endonuclease treatment is shown. Endonuclease treatment destroyed T cell chromosomes with APC telomeres while the total number of APC telomeres remained unaffected. (**c**) Presence of APC-derived telomeres in purified T cell plasma membranes *vs* T cell nuclei 24h after transfer of APC-free supernatants derived as in (**a**). (**d**) Quantification of APC telomere signal after T cell chromatin immunoprecipitation. Telomeres were extracted from Cy-3-PNA labelled APCs with ionomycin then transferred to T cells. The endogenous T cell telomere complex was immunoprecipitated (IP) 24h later with anti-POT1 and the Cy3-fluorescence of APC-derived telomeres quantified with a microplate reader. Control IP, T cell extracts precipitated with irrelevant IgG. Presence of APC-derived telomeres was confirmed by adding DNase directly to the POT1 IP for 10 min at RT prior to fluorescence reading. Two donors. Error bars indicate S.E.M. throughout.

**Extended Data Figure 7.**
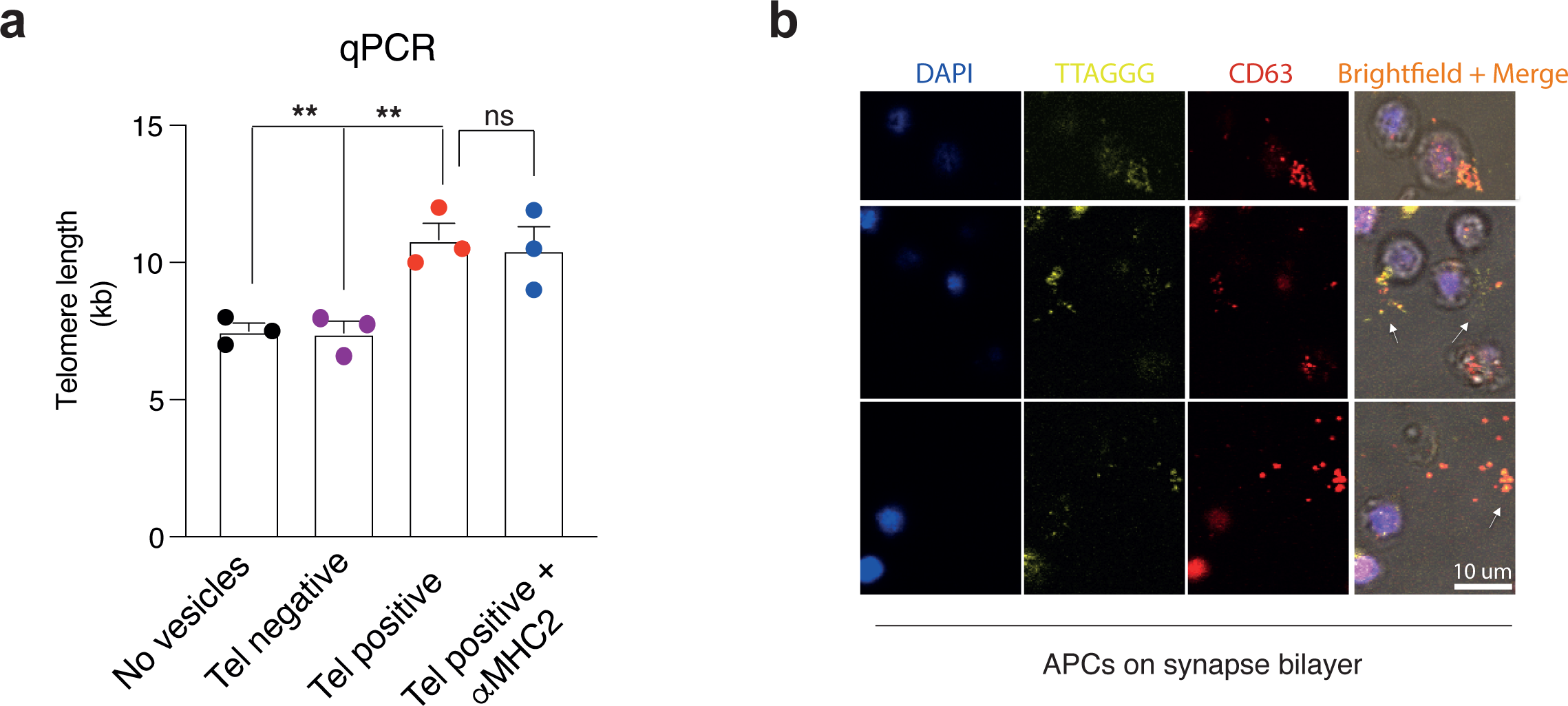
APC-T cell telomere fusion does not require MHC signalling. (**a**) T cells were cultured for 48h with 5,000 telomere vesicles (Tel^+^) purified from APCs by fluorescence activated vesicles sorting, in the presence or absence of blocking MHC II antibodies (1 μg/mL). T cell nuclear extracts were then analysed by qPCR with telomere and single copy reference primers. 5,000 telomere depleted vesicles (Tel^-^) served as control. (**b**) Release of CD63^+^ telomere vesicles on planar bilayers documented by brightfield illumination. Scale bar, 10 μm. Anova with Bonferroni post correction test, \*\**P* < 0.01. ns, not significant. Error bars indicate S.E.M. throughout.

**Extended Data Figure 8.**
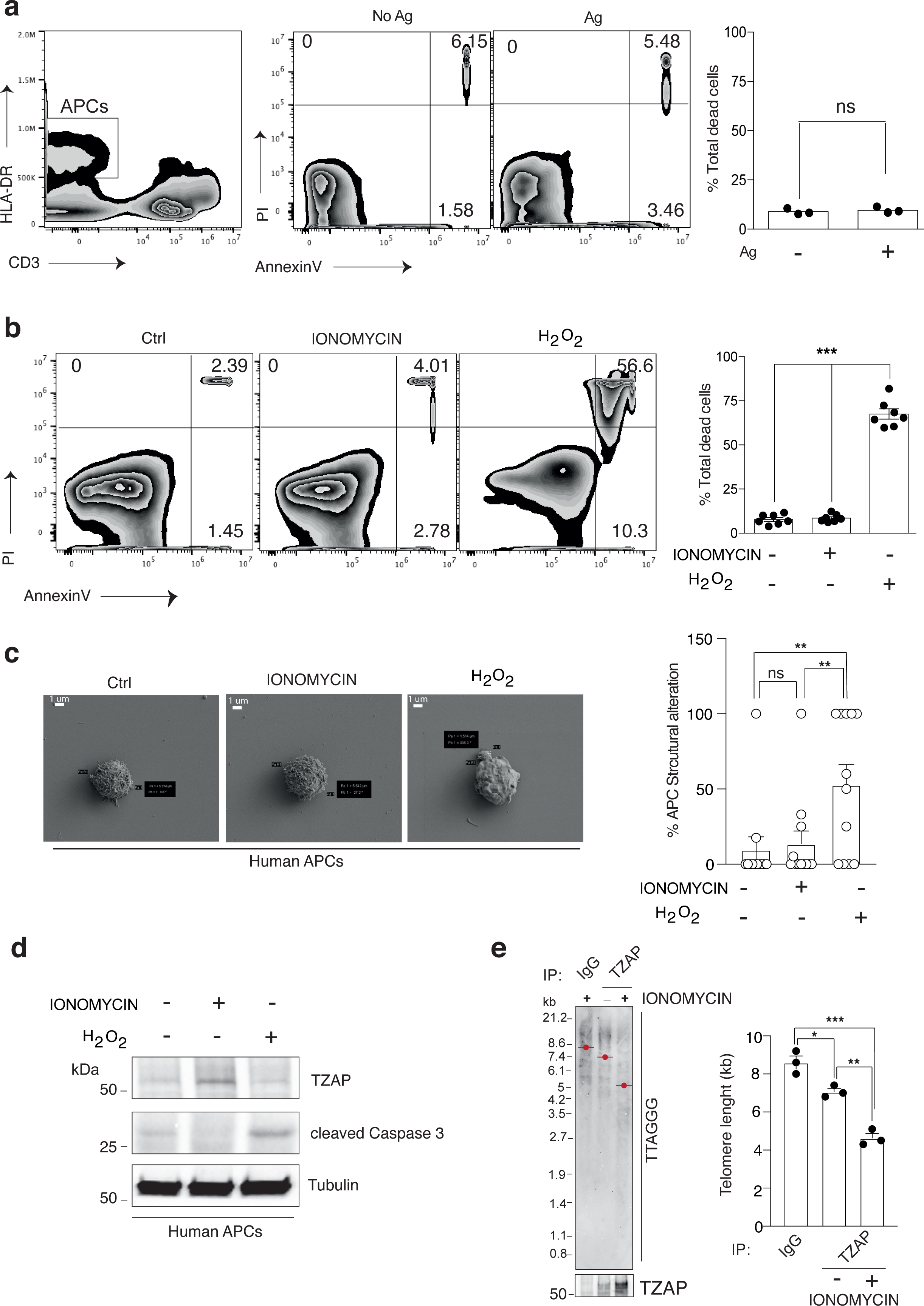
Telomere transfer does not cause APC death. Analysis of cell death of APCs after stimulation with T cell and antigen mix (**a**) or ionomycin (**b**) for 18h. Representative plots (**left**) and percentages (**right**) of total dead cells (Annexin V+/PI-cells, Annexin V+/PI+ and Annexin V-/PI+ cells). These experiments were performed 3 times (**a**) or 7 times (**b**). APCs treated with H2O2 (500μM) served as positive control throughout experiments. (**c, left**) FESEM micrographs (10,000x) of resting APCs or activated APCs upon treatment with ionomycin or H2O2 for 18h. Scale bar 1μm. (**c, right**) Pooled data from 12 micrographs (3-5 APCs per micrograph at 10,000x magnification) depicting % APCs with structural alterations (blebbing or membrane damage). (**d**) APCs treated as in (**b**) were analysed by immunoblot against cleaved-caspase 3 as apoptosis marker, and the telomere trimming factor TZAP. Representative of 3 experiments. Tubulin served as loading control. (**e**) Representative TRF analysis (**left**) and pooled data (n=3; **right**) of TZAP dependent telomere trimming *in vitro*. TZAP was immunoprecipitated from APCs cultured with or without ionomycin for 18h and the immunoprecipitates were incubated for additional 18h at 30°C with genomic DNA extracted from resting APCs. TRF analysis determined telomere shortening upon incubation with TZAP but not control IgG immunoprecipitates that was enhanced by ionomycin. In (**a**), student’s t test, in (**b**, **c, e**), Anova with Bonferroni post correction test, \*\**P* < 0.01. Error bars indicate S.E.M. throughout.

**Extended Data Figure 9.**
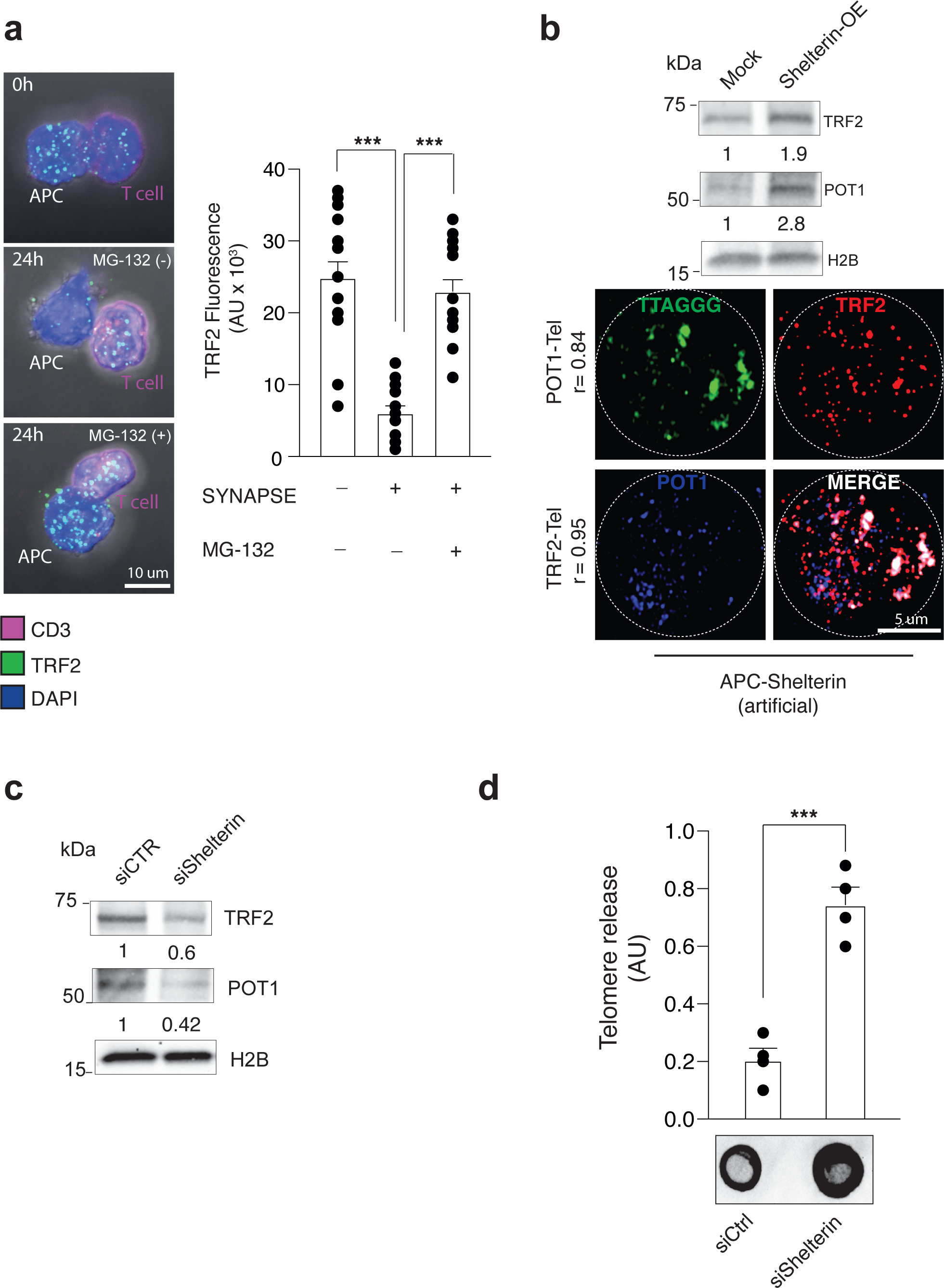
Additional shelterin data. (**a**) Representative images (**left**) and quantification (**right**) of TRF2 levels by immunofluorescence in APCs left either untreated, or pre-incubated with the proteasome inhibitor MG-132 (1μM) then conjugated with T cells for 24h. T=0, initial time at which conjugates are observed (20 min). Images were z-stacked and the raw telomere integrated fluorescence signals (AU, arbitrary units) are shown. 48 conjugates were analysed. Scale bar, 10 μm. (**b, top**) Validation of shelterin overexpression (TRF2 and POT1) by immunoblotting in APCs. Numbers indicate shelterin overexpression efficiency. (**b, bottom**) IF-FISH demonstrating recruitment of artificial shelterin factors (POT1 and TRF2) to telomeres in APCs transduced with lentiviral vectors and activated with ionomycin for 18h. The Pearson’s co-localization score for artificial POT1 and TRF2 with APC telomeres are shown. Scale bar, 5 μm. (**c**) Immunoblot analysis of TRF2 and POT1 following indicated siRNA treatment in human APCs. H2B served as loading control. The numbers indicate knock-down efficiency. Representative of 3 separate experiments. (**d**) Dot-blot analysis of telomeric DNA from the EVs released by APCs transfected as in (**c**) and activated with ionomycin for 18h. Quantification of the arbitrary units (AU) from 4 independent experiments (**top**) and representative dot-blot using TelC probe (**bottom**). In (**a**), Kruskal-Wallis with Dunn’s post-correction test; in (**d**), Mann-Whitney test. \*\**P* < 0.01. Error bars indicate S.E.M. throughout.

**Extended Data Figure 10.**
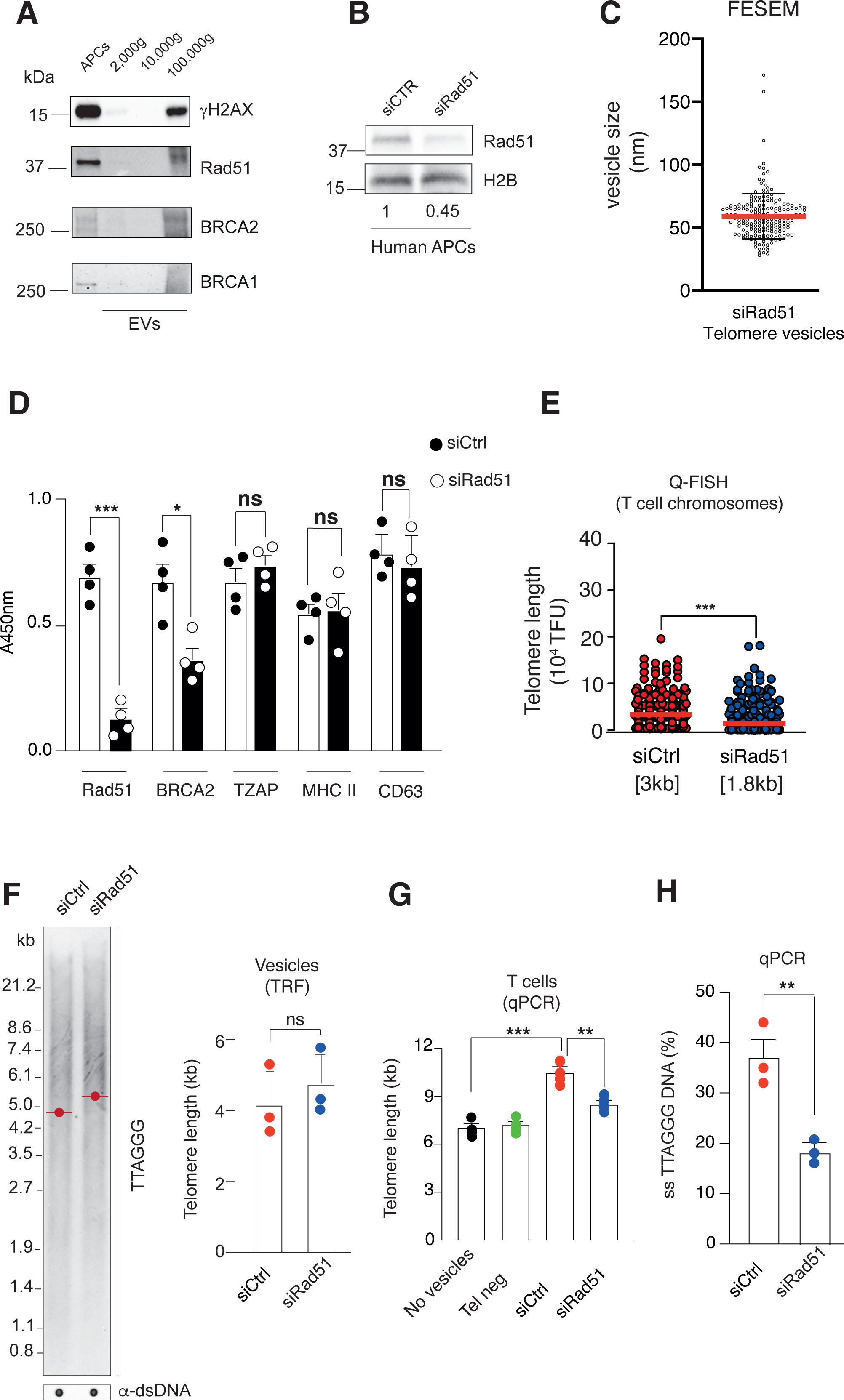
Characterization of Rad51-deficient telomere vesicles. (**a**) Immunoblot analysis of the indicated DNA damage factors in EV pellets derived by sequential centrifugation of APC supernatants following activation with ionomycin for 18h. APCs served as whole cell lysate control. Representative of 3 separate experiments. (**b**) Immunoblots of Rad51 following indicated siRNA treatment in human APCs. H2B served as loading control. The numbers indicate knockdown efficiency. Representative of 3 separate experiments. (**c**) FESEM analysis of 200 siRad51 telomere vesicles purified by fluorescence activated vesicle sorting among cell-free supernatants of APCs transfected as in (**b**) then activated with ionomycin for 18h. (**d**) Protein cargo in siRad51 or siCtrl telomere vesicles by indirect ELISA. Telomere vesicles (250 per sample, in triplicate wells) were purified by fluorescence activated vesicle sorting from cell-free supernatants of APCs transfected with either siCtrl or siRad51 short-interference RNA, then stimulated with ionomycin for 18h. Pooled data from 4 experiments are shown. (**e**) Metaphase Q-FISH coupled to TFL software analysis showing elongation of individual T cell chromosome ends upon transfer of telomere vesicles derived from APCs. Each dot represents an individual T cell chromosome with APC-derived telomeres; 370 (siCtrl) and 378 (siRad51). (**f**) TRF analysis comparison of siCtrl *vs* siRad51 EVs isolated by 100,000g ultracentrifugation following activation with ionomycin for 18h. Pooled data from 3 experiments are shown (**right**). (**g**) Reduced fusion (elongation) between siRad51 telomere vesicles and T cell telomeres. Recipient T cells were treated with 5,000 FACS-purified telomere vesicles (Tel^+^) from siCtrl or siRad51 APCs and total nuclear T cell extracts were analysed by qPCR 48h later. 5,000 telomere depleted vesicles (Tel^-^) were also transferred as control. (**h**) Pooled data (n=3) showing reduced single-strand telomeric DNA in siRad51 *vs* siCtrl telomere vesicles by quantitative amplification of single-stranded DNA (QAOS). In (**d, h**) student’s t test, in Mann-Whitney test, in (**g**) Anova with Bonferroni post-correction test. *p<0.05, **p<0.001 \*\*\**P* < 0.001. Error bars indicate S.E.M. throughout.

**Extended Data Figure 11.**
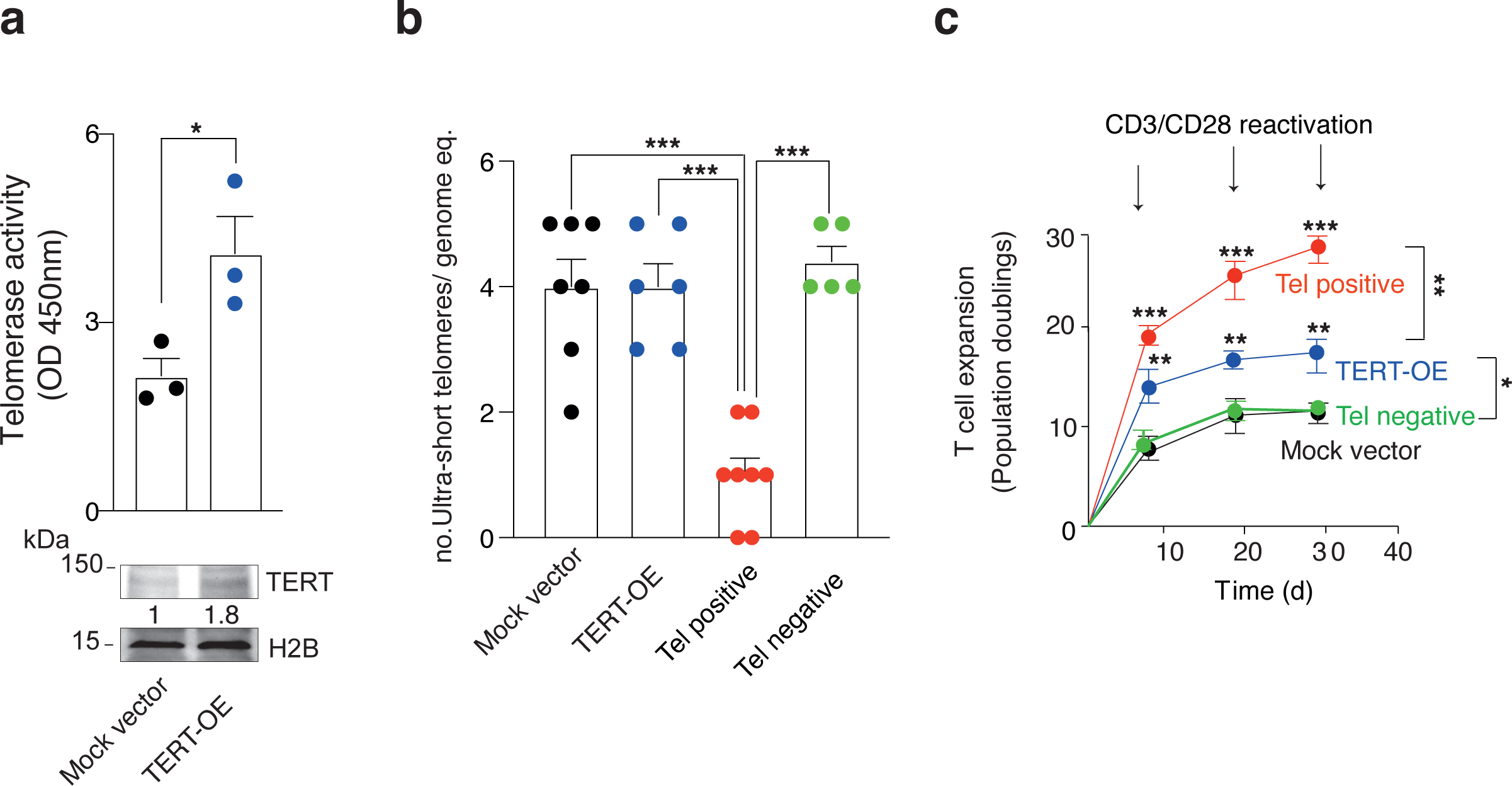
Telomere vesicles, but not telomerase reactivation, eliminate ultra-short telomeres. (**a**) Confirmation of CRISPR-based telomerase enhancement in T cells 96 hours after transduction by TRAP assay (**top**) and immunoblots (**bottom**). The numbers indicate efficiency of CRISPR-based TERT overexpression (TERT-OE). Controls were transduced with mock vectors throughout. **(b)** PCR based assays (U-STELA) showing reduced load of ultra-short telomeres (<3kb) in T cells activated by anti-CD3 plus anti-CD28 for 48h followed by transfer of 1,000 FACS-purified telomere vesicles (Tel^+^) or vector-based overexpression of Tert. Controls were transduced with mock vector or 1,000 telomere depleted vesicles (Tel^-^). Results from 5-8 experiments were normalized to the number of genome equivalent used in PCR amplification reactions (5 genome equivalent, 1 genome equivalent = 8pg DNA). (**c**) Population doublings of T cells treated as in (**b**), then cultured for up to 30 days. In (**a**), student’s t test, in (**b**, **c**) Anova with Bonferroni post-correction test. *p<0.05, **p<0.001 \*\*\**P* < 0.001. Error bars indicate S.E.M. throughout.

**Extended Data Figure 12.**
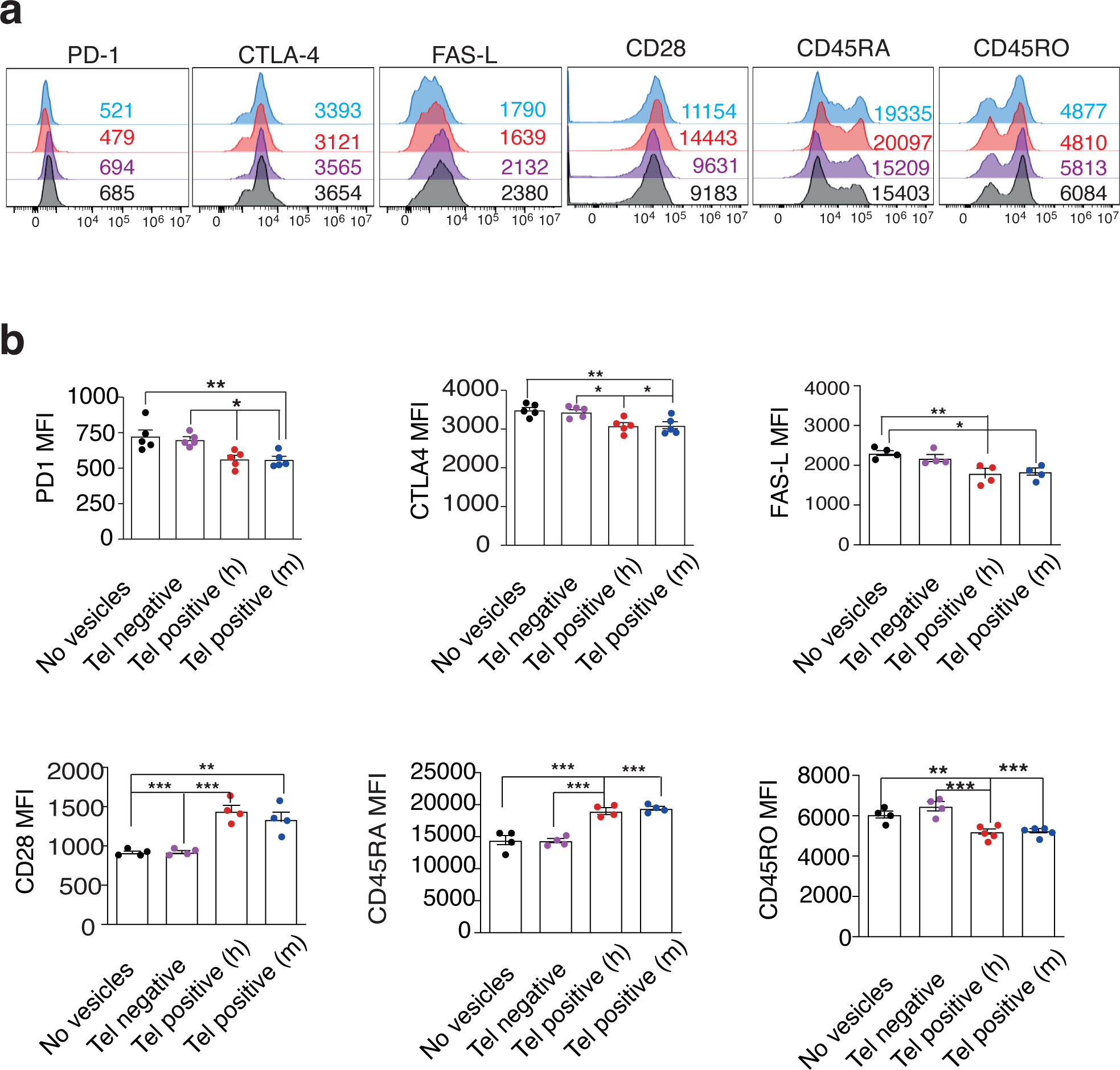
Phenotypic analysis of T cells with APC telomeres. Human T cells were activated with anti-CD3 and IL-2 for 10 days in the presence of 250 FACS-purified telomere vesicles (Tel^+^) derived from either human or mouse APCs prior to multiparametric flow cytometry. Control T cells were activated without any vesicle or with 250 telomere depleted vesicles (Tel^-^) obtained by fluorescence activated vesicle sorting. (**a**) Representative plots and (**b**) pooled data from 5 experiments are shown. In (**b**) Anova with Bonferroni post correction test. *\*P < 0.05; **P< 0.001 and ***P < 0.001*.

**Extended Data Figure 13.**
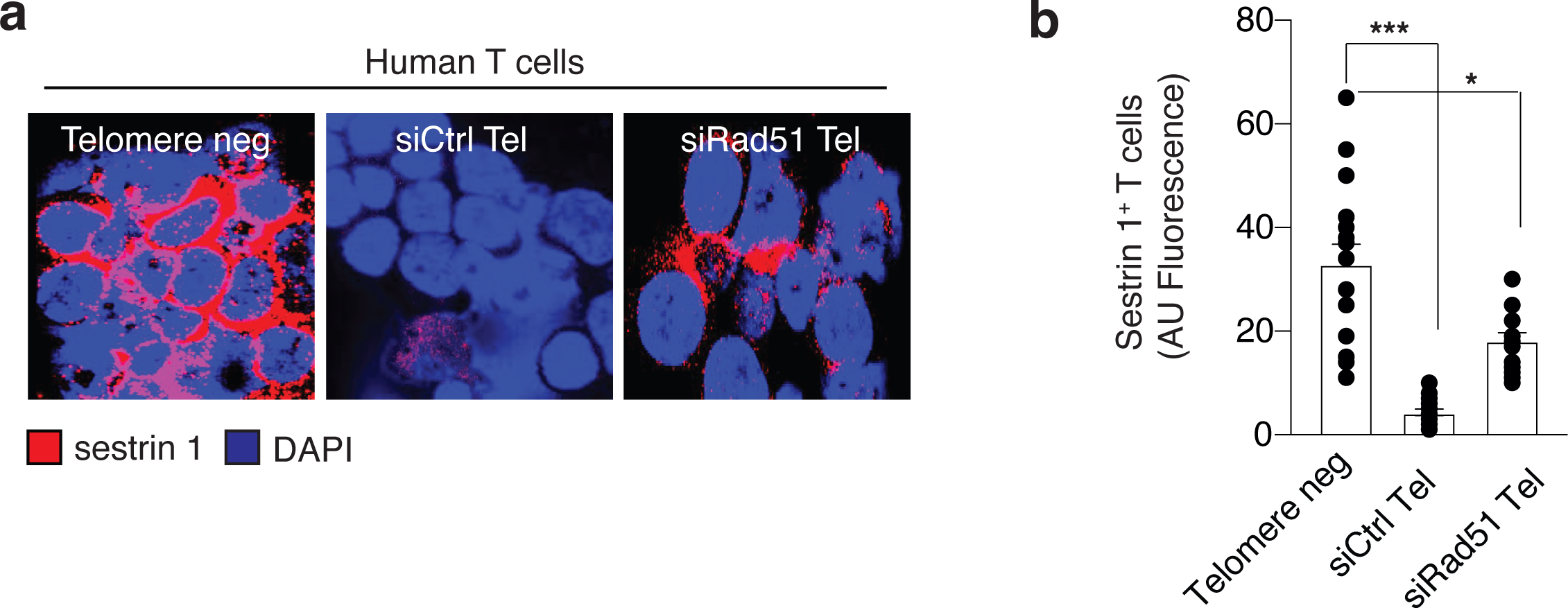
T cells with APC telomeres do not upregulate sestrins. (**a**) Telomere vesicles were extracted with ionomycin from APCs that had been previously transfected with either siCtrl or siRad51 RNAs then transferred to T cells. T cells were then activated by anti-CD3 plus anti-CD28 for 10 days. IF demonstrated that activated T cells do not up-regulate senescence-associated sestrin proteins (sestrin 1) upon transfer of Tel^+^ vesicles. By contrast, T cells up-regulated sestrins (albeit to a lesser extent) upon transfer of siRad51 Tel^+^ vesicles. Two donors. In (**b**), Kruskal-Wallis with Dunn’s post-correction test. *\*P < 0.05; ***P < 0.001*.

**Figure S14.**
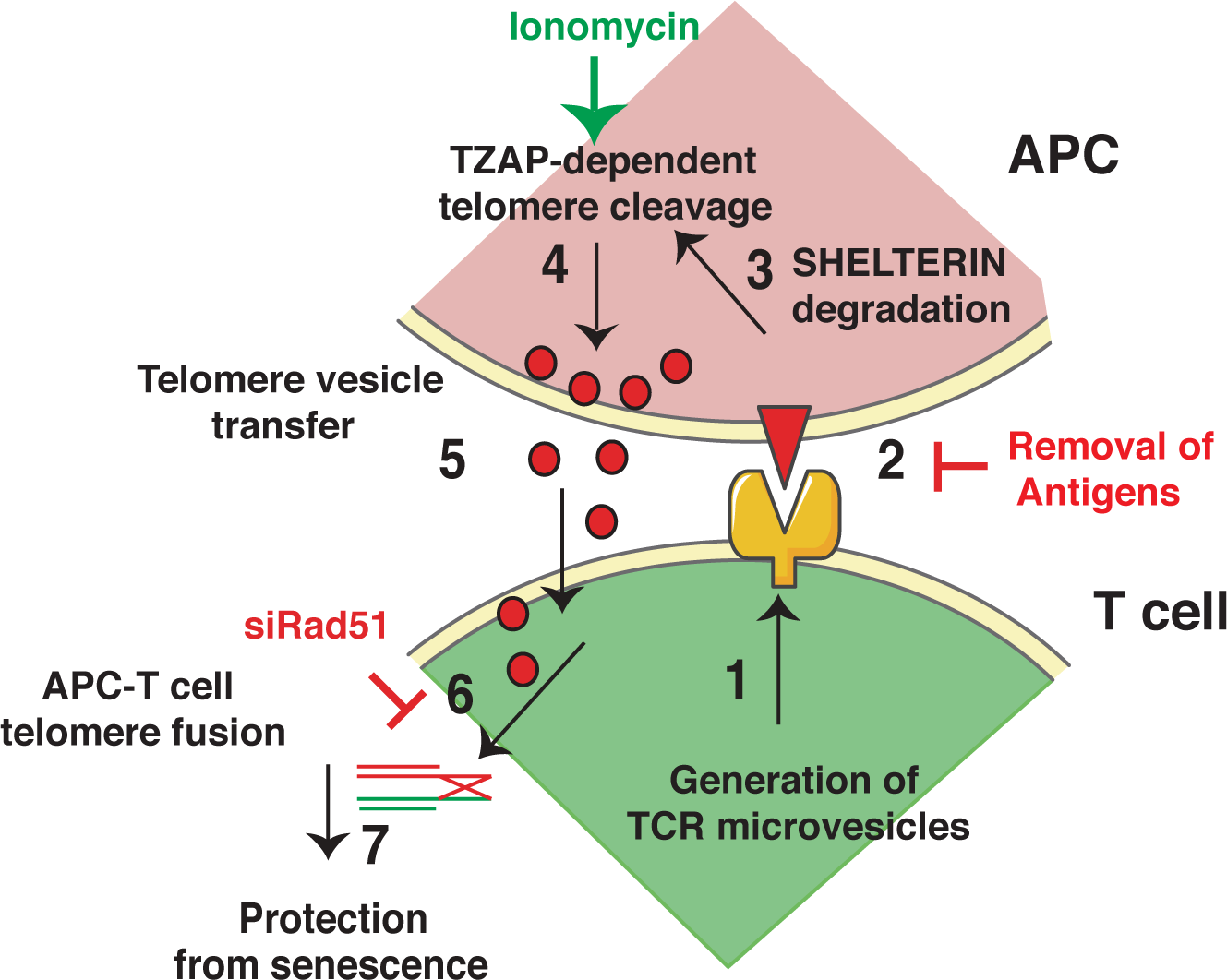
APCs donate telomere vesicles to promote T cell lifespan. Upon antigen-specific contacts with T cells (1-2), APCs degrade shelterin (3) to donate telomeres (4), which are cleaved by TZAP and then transferred in vesicles at the synapse with T cells (5). Telomere vesicles are devoid of shelterin but retain the Rad51 recombination factor that mediates APC-T cell telomere fusion causing an average lengthening of 3000 base pairs (6). The result is a T cell that escapes senescence within 24 hours of synaptic interactions (7). That T cell will become an antigen-specific memory cell.

